# Phonetic underpinnings of sound symbolism across multiple domains of meaning

**DOI:** 10.1101/2024.09.03.610970

**Authors:** Simon Lacey, Kaitlyn L. Matthews, K. Sathian, Lynne C. Nygaard

## Abstract

Sound symbolism occurs when the sound of a word alone can convey its meaning, e.g. ‘balloon’ and ‘spike’ sound rounded and pointed, respectively. Sound-symbolic correspondences are widespread in natural languages, but it is unclear how they are instantiated across different domains of meaning. Here, participants rated auditory pseudowords on opposing scales of seven different sound-symbolic domains: shape (rounded-pointed), texture (hard-soft), weight (light-heavy), size (small-big), brightness (bright-dark), arousal (calming-exciting), and valence (good-bad). Ratings showed cross-domain relationships, some mirroring those between corresponding physical domains, e.g. size and weight ratings were associated, reflecting a physical size-weight relationship, while others involved metaphorical mappings, e.g., bright/dark mapped onto good/bad, respectively. The phonetic features of the pseudowords formed unique sets with characteristic feature weightings for each domain and tended to follow the cross-domain ratings relationships. These results suggest that sound-symbolic correspondences rely on domain-specific patterns of phonetic features, with cross-domain correspondences reflecting physical or metaphorical relationships.

## INTRODUCTION

Sound symbolism refers to instances in which the sound of a word can be enough to convey its meaning (Nuckolls, 1999; Svantesson, 2017); for example, ‘balloon’ and ‘spike’ sound like, and refer to, rounded and pointed objects respectively (Sučević et al., 2015). This idea is at least as old as the Platonic *Cratylus* dialog (Ademollo, 2011), but modern linguistics has taken the opposite view: that the relationship between sound and meaning is almost completely arbitrary (e.g., de Saussure, 1916/2009; Hockett, 1959; Pinker, 1999), and that instances of sound symbolism are too few to be functionally meaningful (see Lev-Ari & McKay, 2023). Alongside this, however, is a large body of work showing that humans are sensitive to sound-symbolic mappings, the classic being the ‘kiki-bouba’ effect, in which pseudowords such as ‘kiki’ or ‘takete’ are reliably associated with pointed shapes, while ‘bouba’ or ‘maluma’ are associated with rounded shapes (Köhler, 1929, 1947; Ramachandran & Hubbard, 2001). Moreover, recent work shows that, far from occurring sporadically, sound symbolism regularly occurs in thousands of languages, across different language families, and even in isolates (a linguistic lineage with a single member) (Blasi et al., 2016). Sound symbolism also appears to have a long linguistic history, for example, occurring among the earliest surviving Sumerian poetic texts from approximately 2500 – 2000 BCE (see Johnson, 2010).

Most studies of sound symbolism investigate a single mapping and by far the most extensively researched sound-symbolic domain is shape; other domains studied include size (Sapir, 1929), brightness (Newman, 1933; Hirata et al., 2011), and gender (Sidhu & Pexman, 2015). Some studies of the sound-to-shape mapping have investigated its phonetic underpinnings, examining the roles of voiced and unvoiced consonants (Cuskley et al., 2017), obstruents and sonorants (McCormick et al., 2015), consonants and vowels (Fort et al., 2015; Nielsen & Rendall, 2011), vowel formants (Knoeferle et al., 2017), and rounded and unrounded vowels (Maurer et al., 2006; McCormick et al., 2015). More recent studies have explored the influence of acoustic parameters derived from spectro-temporal and voice characteristics (Lacey et al., 2020; Akita, 2021; Villegas et al., 2023). However, investigating sound symbolism by reference to a single domain has obvious limitations in that the relative importance of these factors, phonetic or acoustic, might change depending on the particular sound-symbolic mapping. Furthermore, examining multiple domains enables us to consider whether sound-symbolic mappings that differ with respect to phonological structure can nonetheless be grouped by reference to higher-order factors such as intensity or magnitude (Sidhu & Pexman, 2018), arousal (Aryani et al., 2020), or categories such as activity, valence, potency, and novelty (Osgood et al, 1957; Sidhu et al., 2022).

Few studies have investigated more than a single domain of meaning and, in some cases, the number of domains comes at the expense of the number of items. For example, Miron (1961) examined 15 domains but obtained ratings for only 50 pseudowords, while Johansson et al. (2020) investigated 344 basic concepts in representative languages from 245 language families but using only a single word for each concept. Most recently, Sidhu et al. (2022) obtained ratings for 24 domains using a set of only 40 pseudowords. By contrast, Westbury et al. (2018) investigated only 5 domains but used a set of 7996 pseudowords. Tzeng et al. (2017) found that participants could select the correct sound-symbolic meaning for 80 foreign real words covering only 4 domains. Finally, Winter et al. (2017) obtained iconicity ratings for 3001 real words, finding that ratings were higher for words relating to the five sensory domains than for words belonging to more abstract domains. For completeness, Greenberg and Jenkins (1966) investigated 26 domains, but only in relation to individual consonants and vowels (though this is important for comparison to the phonetic features of both real and pseudowords). We should also note that Winter et al. (2023) obtained iconicity ratings for more than 14000 real words, but these were not organized into domains.

Although the studies reviewed above examined sound symbolism in multiple domains, few investigated whether, and how, different domains relate to each other. Factor analyses have shown that sound-symbolic domains can be grouped according to a relatively small number of higher-order factors. For example, an ‘evaluation’ factor, more recently re-labeled ‘valence’ (Sidhu et al., 2022), includes the good/bad and beautiful/ugly domains (Osgood et al., 1957; Miron, 1961). Similarly, a ‘potency’ factor includes heavy/light, large/small, and strong/weak (Osgood et al., 1957; Miron, 1961; Sidhu et al., 2022). A third factor, ‘activity’ or ‘arousal’ (Sidhu et al., 2022) includes active/passive and fast/slow (Osgood et al., 1957), hard/soft (Miron, 1961; Sidhu et al., 2022) and rounded/pointed (Sidhu et al., 2022). However, which factors emerge depends on the selected domains; for example, Sidhu et al. (2022) introduced a new factor, labelled ‘novelty’, whose members included realistic/fantastic and ordinary/unique, among others. Cross-domain relationships were not the focus of prior factor analyses but can be inferred from positive and negative loading. For example, passive/active and tense/relaxed loaded onto the ‘activity’ factor positively and negatively, respectively, (Sidhu et al., 2022) from which we can infer that the underlying scales were negatively correlated, i.e. that passive pseudowords were also rated as relaxed but not tense while the reverse was true for active pseudowords.

Network analyses, a novel way of analyzing cross-domain relationships, were carried out by Sidhu et al. (2022, Figure 5) on ratings of auditory pseudowords. Consistent with the factor analyses reviewed above, these showed strong connections between, for example, the size and weight domains but also between the valence (good/bad) and emotion (happy/sad) domains (Sidhu et al., 2022). Further cross-domain relationships were revealed in a study in which participants listened to sound-symbolic foreign words and chose between their actual meaning, or meanings from other domains, in English (Tzeng et al., 2017). This showed, for instance, that words meaning ‘fast’ also sounded as though they meant ‘pointed’ and ‘moving’, while words meaning ‘big’ and ‘small’ were paired with ‘moving’ and ‘still’, respectively (Tzeng et al., 2017); these cross-domain groupings were attributed to a common, higher-order dimension such as stimulus intensity or magnitude (Tzeng et al., 2017). In contrast to these two studies, Westbury et al. (2018) presented pseudowords visually and only required a simple yes/no decision as to whether each would make a good English word to denote a particular concept. The probability that a pseudoword would be considered to mean ‘large’ was positively correlated with the probability of also being considered to mean ‘round’ and ‘masculine’, and negatively correlated with meaning ‘sharp’ or ‘feminine’ (Westbury et al., 2018).

Here, we collected ratings of a relatively large set of 537 consonant-vowel-consonant-vowel (CVCV) pseudowords (McCormick et al., 2015) for seven meaning domains as a reasonable compromise to the ‘items/domains’ trade-off discussed above. The pseudowords were created using phonemes with reliable, but not exclusive, associations to the shape domain (McCormick et al., 2015; Lacey et al., 2020). We extended our previous work on the shape domain by also obtaining ratings for other perceptual properties of physical objects, i.e., texture, weight, size, and brightness, as well as for the abstract domains of arousal and valence. This choice of domains samples each of the three main factors previously reported, as detailed above. It also enabled us to compare concrete and abstract domains, and different sensory modalities; for example, while brightness is perceived visually, texture is primarily salient to touch (Klatzky et al., 1987), and weight judgments rely on active lifting and associated somatosensory processing (e.g., Valchev et al., 2017). More importantly, examining multiple domains enables us to test a central question in sound symbolism: whether sound-symbolic mappings are domain-specific, in which case the phonetic profile (i.e., the relative contributions of different phonetic features to any particular sound-symbolic mapping) would vary depending on the domain; or whether they reflect mapping to a single, domain-general, abstract factor (such as arousal, magnitude, or valence [see Aryani et al., 2020; Sidhu & Pexman, 2018; Spence, 2011]). In the latter case, the phonetic profile would be similar across domains and reflect that of the hypothesized domain-general factor. For example, if the domain-general factor were arousal then the sound-symbolic phonetic profile for arousal should be preserved across all other domains.

In this paper, we concentrate on the phonetics – the place and manner of articulation of speech sounds – of the pseudowords, that arguably represent an abstract, linguistic characterization of each item. Such phonetic features naturally have acoustic consequences and these are the subject of a companion paper (Lacey et al., 2024). Participants in the present study listened to spoken pseudowords and rated how rounded, good, exciting, etc. each one sounded on a seven-point Likert scale. Cross-domain relationships, for example whether pseudowords rated as small/big were also rated as light/heavy, respectively, were assessed via conventional correlations between the rating scales. The contributions of different phonetic features to each sound-symbolic mapping were assessed via a linear regression separately for each domain.

Our predictions for the phonetic features involved in the sound-symbolic mapping for each domain are grounded in previous findings and limited to the phonetic features used in our pseudoword set (note that this set is not exhaustive of English phonemes or phonetic features, for example it does not include other fricatives like /h/ and /∫/, the glides /w/ and /y/, nor low vowels). Full details of the consonants and vowels used in the pseudoword set, and their manner, voicing, and place of articulation, are provided in Table 1.

**Table 1:**
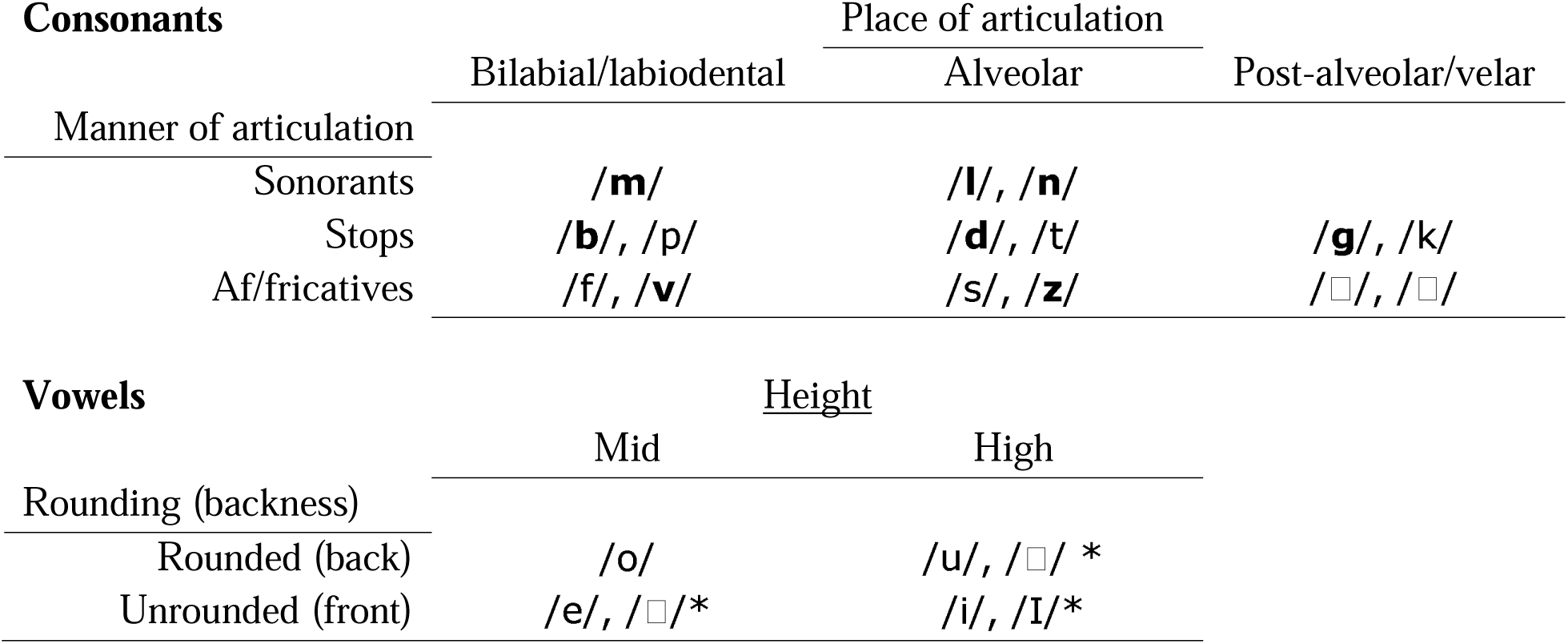
Phonetic and articulatory features of consonants and vowels used in creating the CVCV pseudowords. Voiced consonants are in **bold** type; vowels marked * only occur in the first vowel position as they do not occur in word-final positions in English.

For the shape domain, roundedness is associated with voiced (e.g., /b/, /d/) rather than unvoiced (e.g., /t/, /k/) consonants, and with back/rounded vowels (e.g., /u/ or /o/) (Köhler, 1929, 1947; Ramachandran & Hubbard, 2001; McCormick et al., 2015), while pointedness is associated with stops (e.g., /p/ and /t/) rather than sonorants (e.g., /m/ or /l/), and with front/unrounded vowels (e.g., /i/ or /e/: Köhler, 1929, 1947; Ramachandran & Hubbard, 2001; McCormick et al., 2015). Consistent with these studies, Westbury et al. (2018) reported that sharpness (which we take to be synonymous with pointedness since it was paired antonymically with roundness rather than bluntness) was associated with velar consonants e.g., /g/ or /k/, while roundedness was associated with nasal consonants (Westbury et al., 2018: of the three nasal consonants in English, the present study uses /m/ and /n/; the other is /ŋ/ as in ‘sing’ [Reetz & Jongman, 2020]). A more recent study generally confirms the findings for consonants, with roundedness associated with sonorants and voiced stops, while pointedness was associated with unvoiced stops and voiced fricatives (Sidhu et al., 2022).

The classic finding for sound-symbolic size associations is that front and back vowels reflect small and large size, respectively (Sapir, 1929; replications can be found in Auracher, 2017; Sidhu et al., 2022). Among the consonants, fricatives are associated with small size, and stops with large size (Preziosi & Coane, 2017). The associations for fricatives may depend on voicing, with unvoiced/voiced fricatives associated with small and big, respectively (Sidhu et al., 2022). Similarly, while Westbury et al. (2018) found that large size was associated with voiced consonants, Sidhu et al. (2022) reported that this only related to voiced stops. Moreover, sonorants – which are always voiced – are associated with small size (Sidhu et al., 2022).

For the hard/soft dimension of texture, hardness is associated with the alveolar affricate /□/ and the velar stop /k/ when textures were directly perceived via touch (Sakamoto & Watanabe, 2018) and with voiced fricatives and unvoiced stops generally (Sidhu et al., 2022; see also Greenberg & Jenkins, 1966). By contrast, softness is associated with the bilabial stops /b/ and /p/, the alveolar sonorant /n/ for felt textures (Sakamoto & Watanabe, 2018) and with sonorants generally (Sidhu et al., 2022; Greenberg & Jenkins, 1966). However, there appear to be no specific associations to vowels (Sidhu et al., 2022).

For the weight domain, stops were described as heavy regardless of voicing (Greenberg & Jenkins, 1966). However, in another study heaviness was associated with voiced stops and voiced fricatives, while lightness was associated with sonorants and unvoiced fricatives (Sidhu et al., 2022). Front vowels have been associated with lightness, and back vowels with heaviness (Walker & Parameswaran, 2019; Sidhu et al., 2022).

Brightness is associated with unvoiced stops /k/ and fricatives /s/, and front vowels /i/, /I/, while darkness is associated with voiced stops /b/ /g/and sonorants /m/, and back vowels /u/ /o/ (Newman, 1933). These findings were only partially replicated by Greenberg & Jenkins (1966) where the darkest consonants consisted of both voiced and unvoiced stops, while the brightest consonants consisted of both voiced and unvoiced fricatives, and sonorants.

For the arousal domain, sonorants and back vowels are associated with pseudowords perceived as calming while exciting pseudowords were associated with voiced fricatives, unvoiced stops, and front vowels (Sidhu et al., 2022). In a study in which ‘kiki’-like pseudowords were considered more arousing than ‘bouba’-like pseudowords (Aryani et al., 2020), unvoiced consonants, sibilants, and short vowels were associated with more arousing pseudowords and voiced consonants and long vowels were associated with less arousing pseudowords (Aryani et al., 2018)

Sound-symbolic valence is relatively unexplored and might depend on the definition: moral good/bad vs emotional happy/sad, for example; here, we chose the good/bad dimension. For real words across several languages, longer/shorter initial phonemes predicted positive/negative valence, respectively (Adelman et al., 2018), which suggests that sonorants and obstruents will be associated with good/bad, respectively. In a study of names for the complex concept of villains (Uno et al., 2020), voiced obstruents (stops, affricates, and fricatives) were associated with negative valence. A recent study of constructed, fictional, languages, intended by their creators to sound either harsh and evil (e.g., Klingon, Orkish) or pleasant and good (e.g., Sindarin, Quenya), suggests that voiced speech sounds are perceived more positively than unvoiced speech sounds (Mooshammer et al., 2023). Consistent with this finding, unvoiced consonants, sibilants, and short vowels have been associated with negative valence, while voiced consonants and long vowels were associated with positive valence (Aryani et al., 2018). The front vowel /i/ and the back vowel /o/ are associated with positive and negative valence respectively (Körner & Rummer, 2022). Note that Sidhu et al. (2022) found no significant effect of either consonants or vowels for their pseudowords.

Across domains, we test whether different sound-symbolic mappings reflect a single domain-general factor or multiple domain-specific factors. For example, if all sound-symbolic mappings can be traced to a general factor, such as arousal, then every domain should have a phonetic profile similar to that for arousal (as described above). Alternatively, finding that each domain has a unique phonetic profile would be evidence in favor of a domain-specific account of sound symbolism.

## METHODS

### Participants

Participants were recruited, and compensated for their time, online via the Prolific participant pool (https://prolific.co: see Peer et al., 2017). The eligibility criteria were that participants be aged between 18 and 35, be native speakers of American English, and have no language or hearing disorders. A total of 646 people took part, but data from 257 were excluded: 73 because they failed a headphone screen (see General Procedures below) and a further 184 because post-task questionnaires (see General Procedures below) indicated that they were bilingual or spoke a second language (120); they used less than 5 values on the full 7-point scale (47); there was a malfunction in stimulus presentation (8); they experienced synesthesia (3), which has been associated with increased sensitivity to sound-symbolic crossmodal correspondences (Lacey et al., 2016); and 6 for other reasons. Thus, the final sample comprised 389 participants (171 male, 208 female, 6 non-binary, 2 agender, 1 gender-fluid, and 1 declined to state their gender; mean age 27 years, 1 month (standard deviation = 8 months). The rating experiments were hosted on the Gorilla platform (https://gorilla.sc: Anwyl-Irvine et al., 2020) where participants also gave informed consent. All procedures were approved by the Emory University Institutional Review Board.

### Auditory Pseudowords

We used the 537 two-syllable CVCV pseudoword set created by McCormick et al. (2015), including only phonemes and combinations of phonemes that occur in the English language, and excluding items that were considered to be homophones of real words. Consonants were sampled from sonorants, affricates/fricatives (henceforth ‘af/fricatives’), and stops; for the af/fricatives and stops, half were voiced and half were unvoiced; vowels were either back/rounded or front/unrounded. Full details of the consonants and vowels, organized by their manner and place of articulation, are provided in Table 1. Because of the high number of possible phoneme combinations, the pseudoword set was restricted to a subset of phonemes and phoneme combinations that were hypothesized, based on prior studies, to reliably map to the rounded/pointed dimension of shape, which was the focus of the original study (see McCormick et al., 2015, for details). Within a given pseudoword, consonants were either both unvoiced (as in /kupo/) or both voiced (as in /gubo/). Pseudowords contained either back/rounded or front/unrounded vowels and did not contain repeated syllables, so possible pseudowords such as /kiki/ and /lolo/ were excluded. For a complete description of the stimulus set, see McCormick et al. (2015); the complete set of pseudowords is available at https://osf.io/ekpgh/.

The pseudowords were recorded in random order, and with neutral intonation, by a female native speaker of American English. Recordings took place in a sound-attenuated room, using a Zoom 2 Cardioid microphone, and were digitized at a 44.1 kHz sampling rate. Two independent judges assessed whether each item was spoken with neutral intonation, accurately produced the intended phonemic content, and sounded consistent with other items (e.g., did not sound faster/slower or louder/softer than other items). For those pseudowords where one of the judges noted that the recording did not conform to any one of these requirements, that item was re-recorded and judged again. Each pseudoword was then down-sampled at 22.05 kHz, which is a standard sampling rate for speech, and amplitude-normalized using PRAAT speech analysis software (Boersma & Weenink, 2012). The pseudowords had a mean (± standard deviation) duration of 457 ± 62 ms.

### General procedures

Participants recruited via Prolific followed a link that took them to the experiment on the Gorilla platform. At the landing page, they could read the consent form and a short description of the task, before clicking on ‘yes’ to continue and complete the rating task, or ‘no’ to exit. Having consented, and before continuing with the rating task, participants completed a headphone check designed to ensure compliance with headphone use for web-based experiments (Woods et al., 2017). Participants who failed the headphone check could take it a second time but if they failed again, while they could still participate, in order to avoid discriminating against those who did not have access to headphones, their data were excluded from analysis. The randomizer function in Gorilla then assigned participants (including those who failed the headphone check) to one of the two opposing rating scales for the relevant domain. Once the rating task had been completed, participants were directed to a series of short questionnaires that asked about any task strategies that had been used, demographic information, language experience and ability, and musical experience and ability. Participants could then read a debriefing statement and exit the experiment. Once participants had completed any one scale, they were excluded from further participation in the study. Thus, the fourteen scales were completed by independent groups of participants.

### Perceptual rating tasks

Participants were randomly assigned to one of fourteen 7-point Likert-type scales and rated all 537 pseudowords on the scale to which they were assigned. The scales represented categorical opposites across seven different meaning domains: shape (rounded/pointed, N =30/30), texture (hard/soft, N = 27/27), weight (light/heavy, N = 29/26), size (small/big, N = 32/31), brightness (bright/dark, N = 24/26), arousal (calming/exciting, N = 30/28), and valence (good/bad, N = 26/23). In order to avoid response bias, one of the scales rated one end of the meaning domain, for example, roundedness, from 1 (not rounded) to 7 (very rounded) and the other rated the other end, i.e., pointedness, from 1 (not pointed) to 7 (very pointed). This also avoided assumptions about contrastive meaning for each domain, i.e., that ‘rounded’ inherently also means ‘not pointed’, or that ‘not exciting’ also means ‘calming’ as opposed to ‘dull’. The instructions included several related terms for the categorical opposites in each domain (see Supplementary Table 1). The 7-point rating scale appeared in the center of the screen on each trial and always listed 1-7 from left to right. Each pseudoword was presented only once, in random order.

### Sample size

In order to get a reliable estimate of the mean rating for each pseudoword, we expected that a sample size of 25-30 participants per scale would be sufficient. McCormick et al. (2015), whose ratings data we used in our previous study (Lacey et al., 2020), had sample sizes of 15 and 16 for their rounded and pointed scales respectively, while other studies included as many as 40+ or 70+ participants (e.g., Miron, 1961). Motamedi et al. (2019) suggest that, for most measures, 10 ratings per item is sufficient, based on simulations. However, the latter largely examined studies using iconicity scales; that is, participants rated how iconic an item was for a suggested meaning on scales that ran, for example, from ‘very opposite’, through ‘arbitrary’, to ‘very iconic’ (Perlman & Lupyan, 2018). But such scales are quite different from the more direct approach adopted here and in other studies, in which participants are asked to rate items on a scale that reflects the meaning itself, e.g., from ‘not pointed’ to very ‘pointed’. The sample size for the correlations between the ratings scales is fixed by the number of pseudowords at 537 and this gives statistical power of >80% to detect a Pearson correlation of .3 when α = .05 (power analysis was carried out in SPSS v29.0 (IBM Corp, Armonk NY).

### Phonetic analysis

We used linear regression to assess the extent to which each phonetic feature was associated with judgments across the different meaning domains. The analysis was conducted in SPSS. All predictors were entered simultaneously, i.e., forced-entry (Field, 2018), meaning that there were no *a priori* assumptions about the relative importance of each of the phonetic features. This approach allowed for predictors to be differently weighted depending on the domain. As detailed below, we modeled manner of articulation, voicing, and place of articulation. The place of articulation for consonants was grouped into bilabial/labiodental, alveolar, and post-alveolar/velar; for vowels, place of articulation was divided into middle and high vowel height (Table 1). Note that grouping the consonants into bilabial/labiodental and post-alveolar/velar groups avoids multicollinearity issues arising from treating similar places of articulation separately.

A regression model was run separately for each of the seven domains, in which each phonetic predictor was evaluated for its influence on perceptual ratings, relative to a reference predictor. For the manner of articulation of consonants, sonorants were the reference predictor for comparison with stops and af/fricatives (collectively referred to as obstruents). Voiced consonants were the reference for unvoiced consonants. For the place of articulation of consonants, bilabial/labiodental consonants were the reference for comparison with alveolar and post-alveolar/velar consonants (each sub-divided into first or second consonant position). For the place/manner of articulation of vowels, front unrounded vowels were the referent for back rounded vowels, and middle height vowels were the referent for high vowels (also sub-divided into first or second vowel position). Positive beta coefficients for the comparison predictors (for example, stops and af/fricatives), would mean that these account for more of the variance than the reference predictor (in this case, sonorants). By contrast, negative beta coefficients would mean that the reference predictor contributes more (in this case, sonorants would account for more of the variance than either stops or af/fricatives).

## RESULTS

### Ratings analysis

Skewness values ranged from -.1 to .7 for the fourteen rating scales, while kurtosis values ranged from -.5 to .2. These values are within the acceptable range of ± 1 for skewness and ± 2 for kurtosis (Hair et al., 2022) and we therefore considered the ratings for each scale to be normally distributed. Within each domain, the two scales were intended to be in opposition to each other, e.g. good vs bad, and therefore were predicted to be strongly negatively correlated. This was, in fact, the case (see the bolded values on the diagonal in Table 2). Thus, for each domain, the two scales adequately reflected the categorical opposites and the meaning domains likely consisted of continuous, contrastive, dimensions.

**Table 2:** Ratings correlations within and between domains. Within each domain, the two scales were significantly negatively correlated (shown in **bold**), indicating that they adequately reflected the opposing dimensions. Boxes indicate contrastive ‘double dissociation’ relationships across domains. Degrees of freedom = 535 in all cases; * p < .05, ** p < .01, *** p < .001; cross-domain comparisons are Bonferroni-corrected for four tests (α = .0125)

|  |  | Shape |  | Texture |  | Weight |  | Size |  | Brightness |  | Arousal |  | Valence |  |
| --- | --- | --- | --- | --- | --- | --- | --- | --- | --- | --- | --- | --- | --- | --- | --- |
|  |  | Rounded | Pointed | Hard | Soft | Light | Heavy | Small | Big | Bright | Dark | Calming | Exciting | Good | Bad |
| **Shape** | Rounded |  |  |  |  |  |  |  |  |  |  |  |  |  |  |
|  | Pointed | **-.32\*\*\*** |  |  |  |  |  |  |  |  |  |  |  |  |  |
| **Texture** | Hard | -.22*** | .78*** |  |  |  |  |  |  |  |  |  |  |  |  |
|  | Soft | .26*** | -.53*** | **-.59\*\*\*** |  |  |  |  |  |  |  |  |  |  |  |
| **Weight** | Light | -.01 | -.50*** | -.61*** | .47*** |  |  |  |  |  |  |  |  |  |  |
|  | Heavy | .01 | .50*** | .65*** | -.49*** | **-.64\*\*\*** |  |  |  |  |  |  |  |  |  |
| **Size** | Small | .06 | .03 | -.01 | .22*** | .26*** | -.20*** |  |  |  |  |  |  |  |  |
|  | Big | .19*** | .24*** | .33*** | -.31*** | -.48*** | .49*** | **-.43\*\*\*** |  |  |  |  |  |  |  |
| **Brightness** | Bright | -.66*** | .35*** | .21*** | -.28*** | -.02 | .01 | -.05 | -.06 |  |  |  |  |  |  |
|  | Dark | .46*** | -.23*** | -.07 | .04 | -.23*** | .17*** | -.20*** | .25*** | **-.53\*\*\*** |  |  |  |  |  |
| **Arousal** | Calming | .36*** | -.67*** | -.66*** | .53*** | .48*** | -.49*** | .10 | -.29*** | -.31*** | .12** |  |  |  |  |
|  | Exciting | -.49*** | .40*** | .23*** | -.23*** | -.01 | .05 | .01 | -.03 | .50*** | -.44*** | **-.33\*\*\*** |  |  |  |
| **Valence** | Good | -.11 | -.32*** | -.33*** | .14** | .39*** | -.39*** | .11* | -.34*** | .17*** | -.30*** | .35*** | .1 |  |  |
|  | Bad | .07 | .31*** | .29*** | -.12** | -.26*** | .28*** | .02 | .21*** | -.08 | .16*** | -.36*** | -.02 | **-.51\*\*\*** |  |

Across domains, there were some ‘double dissociation’ effects in that ratings on the two scales of a given dimension showed opposing correlations with those of another dimension (see boxes in Table 2). Interestingly, these seemed to cluster into cross-domain relationships that reflect either physical or metaphorical associations. For example, for the size and weight domains, pseudowords rated as small were also rated as light but not heavy, while pseudowords rated as big were also rated as heavy but not light. Other instances illustrate more metaphorical relationships. For the valence and weight domains, for example, pseudowords rated good were also positively correlated with light but negatively with heavy, while those rated as bad were positively correlated with heavy but negatively with good. We return to these cross-domain relationships in more detail in the Discussion.

### Phonetic analysis and discussion by domain

In order to evaluate the robustness of the regression results, we first ran several ‘safety’ checks. Firstly, we addressed the problem of multicollinearity, the extent to which two or more predictors are correlated with each other: the stronger such correlations are, the more difficult it becomes to estimate the importance of any predictor on its own. However, following Menard (1995), there were no tolerance values below .2, with observed values ranging from .39 to .98, and variance inflation factors (VIF) ranged from 1.02 to 2.55, well below the threshold value of 10 recommended by Myers (1990: Supplementary Table 2). In addition, the mean VIF of 1.5 was close to 1, as recommended by Bowerman & O’Connell (1990). Thus, we concluded that multicollinearity was not a problem. Secondly, we tested the assumption of independent errors, i.e. the extent to which residual errors are correlated, via the Durbin-Watson test. Field (2018) recommends a conservative approach in which values < 1 or > 3 would be cause for concern. Since our observed values ranged from 1.79 to 2.23 (Supplementary Table 3), we concluded that the assumption of independent errors was met.

Given the number of domains and predictors of the various ratings, we report only the R^2^ and standardized beta coefficients in Table 3; the full regression results are provided in Supplementary Table 4. Wherever possible, the examples given below of the associated phonetic features are drawn from the ten most highly rated pseudowords on each of the two scales for each domain (Table 4); where they are not, the ordinal position of the item is specified. The domains are reported, in approximately descending order of the overall model fit, by reference to the highest R^2^ value for either of the two scales involved.

**Table 3:**
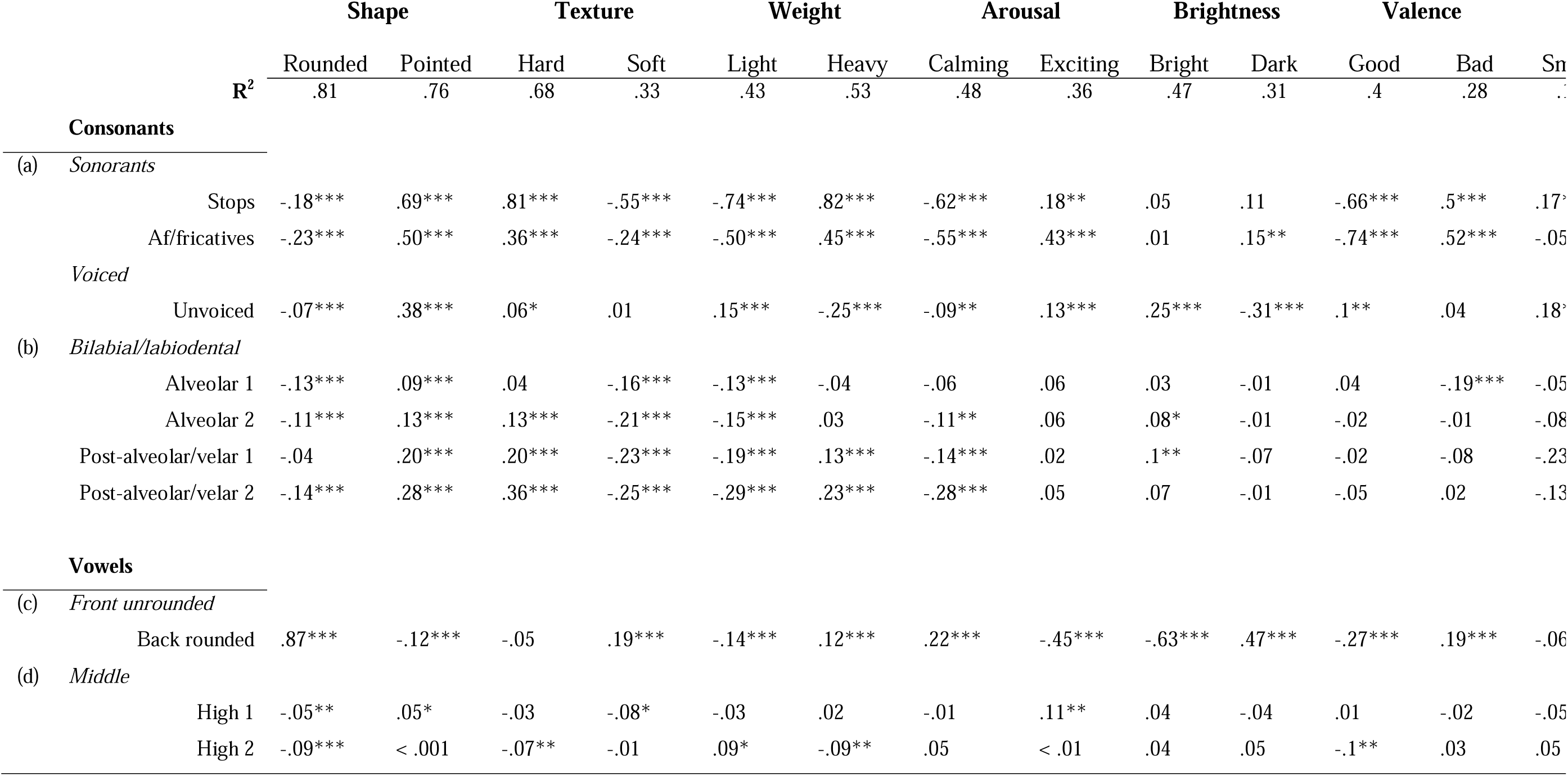
R^2^ and standardized beta coefficients for the regression analysis of the phonetic features for each domain: the dummy-coded reference predictors are italicized; (a) manner and (b) place of articulation for consonants; (c) manner and (d) place of articulation for vowels; 1,2, first or second consonant or vowel position; p values as for Table 2.

**Table 4:**
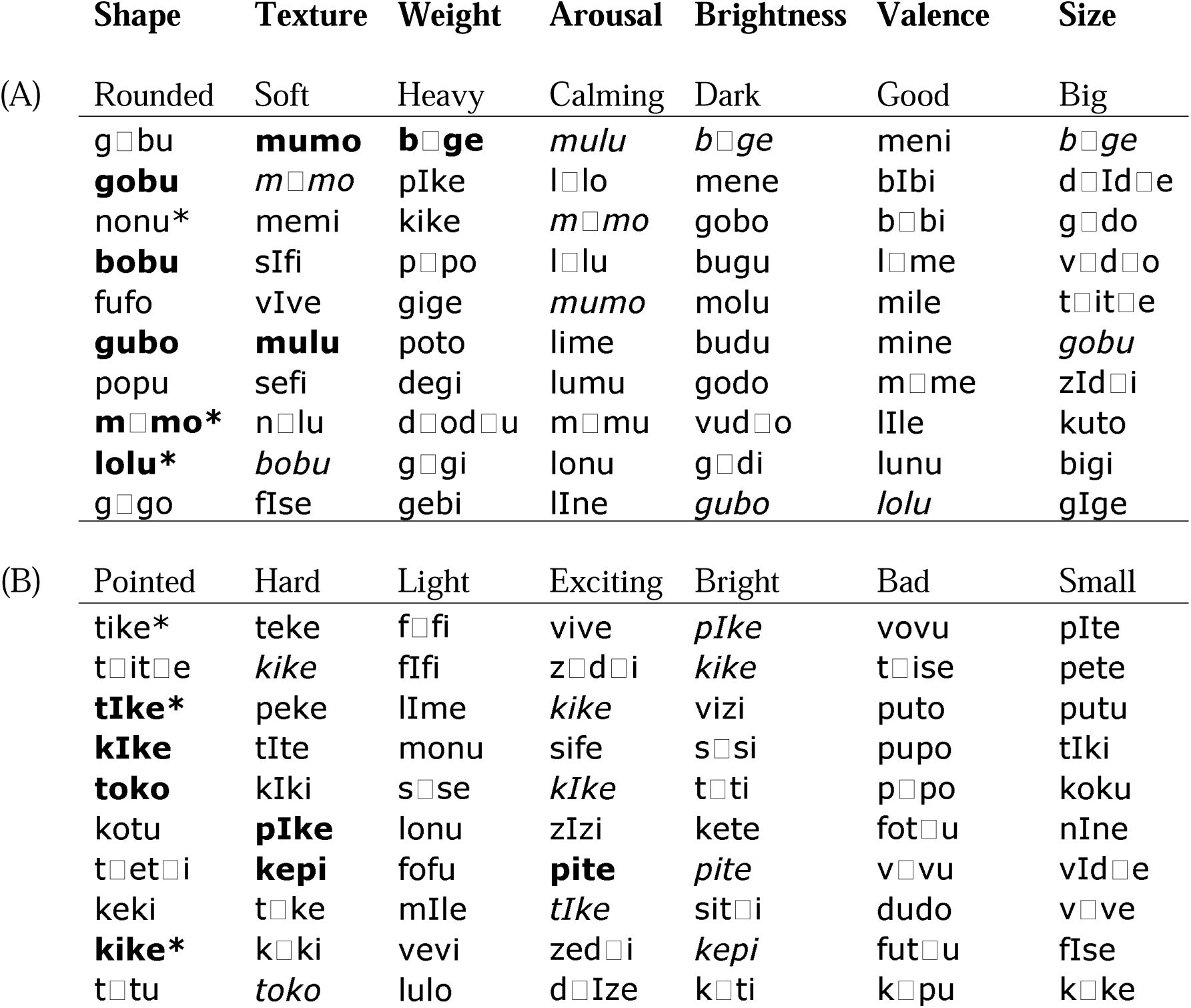
Repetitions among the 10 most highly-rated pseudowords for each scale in each domain. Scales are grouped by phonetic and acoustic (spectro-temporal) similarity (see Lacey et al., 2023). Bold indicates pseudowords that repeat within the group and italics indicate the repetitions of these; most pseudowords were unique in each group – A: 52 (74.3%); B: 54 (77.1%). Shape pseudowords marked * were also in the top 10 for the shape ratings obtained by McCormick et al. (2015).

#### Shape

Unsurprisingly, since the pseudoword set was originally designed to investigate sound-to-shape mappings, the shape domain had the best model fit between phonetic features and ratings, with R^2^ of .81 for the rounded scale and .76 for the pointed scale. Consistent with previous studies (Köhler, 1929, 1947; Ramachandran & Hubbard, 2011; McCormick et al., 2015), roundedness was predicted better by sonorants than obstruents, for example, /m/ and /n/ in /m□mo/ and /nonu/. There was a small but significant advantage for voiced over unvoiced consonants, exemplified by /b/ and /g/ in /gobu/. In terms of place of articulation, bilabial and labiodental consonants were good predictors of roundedness, with /b/, /m/, and /p/ all appearing in the most highly rated rounded pseudowords. Back rounded vowels were strongly associated with roundedness (consistent with Köhler, 1929, 1947; Ramachandran & Hubbard, 2011; McCormick et al., 2015; Sidhu et al., 2022), for example, /o/ and /□/ in /m□mo/, and there was a small but significant effect for mid-height vowels, such as /o/ in /lolu/, over high vowels. By contrast, ratings of pseudowords as pointed were more associated with stops and af/fricatives (consistent with Sidhu et al., 2022), such as /t/ and /□/ respectively, in /tike/ and /□i□e/; unvoiced consonants, such as /t/ and /k/ in /t□tu/ and /kike/, were more predictive of pointedness than voiced consonants (replicating McCormick et al., 2015). The alveolar and post-alveolar/velar consonants were strongly associated with pointedness in either consonant position, exemplified by /t/ and /k/ respectively. Vowels associated with pointedness were predominantly front unrounded vowels, for example /I/, /i/ and /□/ in /kIke/ and /t□ti/, the 30^th^ most pointed pseudoword, (consistent with Köhler, 1929, 1947; Ramachandran & Hubbard, 2001; McCormick et al., 2015; Sidhu et al., 2022). In addition, there was a significant prevalence for high vowels (/i/ and /I/) in the first vowel position of the CVCV pseudowords, for example, /tike/ and /tIke/.

#### Texture

In the texture domain, although R^2^ for the hard scale was .68 (the third highest value overall), the model fit was the most asymmetrical for any domain, in that R^2^ for the soft scale was only .33 (the fourth lowest value overall). Pseudowords were more likely to be rated as sounding hard if they contained stops and af/fricatives rather than sonorants (Greenberg & Jenkins, 1966; Sakamoto & Watanabe, 2018; Sidhu et al., 2022), for example the stops /p/ and /k/ in /peke/ and /toko/. Although the af/fricatives did not occur in the 30 hardest-rated items, this likely reflects the relative weighting of stops and af/fricatives, .81 and .36 respectively. There was a small but significant influence of unvoiced consonants for hardness ratings (Sidhu et al., 2022), for example /t/ and /p/ in /teke/ and /pIke/. In terms of the place of articulation, hardness ratings were associated with alveolar consonants in the second position, such as /t/ in /toko/, and with the post-alveolar consonants in either position, for example, /k/ and /g/ in /kike/ and /degi/, the 13^th^ hardest pseudoword. There was no significant influence of vowel roundedness but there was a small effect for vowel height. Pseudoword softness, meanwhile, was associated with sonorants (see also Sidhu et al., 2022; Greenberg & Jenkins, 1966), such as /m/, /n/, and /l/ in /mumo/ and /n□lu/. In terms of place of articulation, labiodental consonants such as /f/ in /sefi/ also predicted softness, as did back rounded vowels, for example, in /mulu/.

#### Weight

The next best model fit was for the weight domain, with R^2^ of .43 for the light scale and .53 for the heavy scale. Pseudowords rated as light were associated with sonorants (as in Sidhu et al., 2022) such as /l/ and /n/ in /lonu/, with a preference for unvoiced over voiced consonants, for example, /f/ and /s/ in /f□fi/ and /sεse/ (consistent with the association with unvoiced fricatives in Sidhu et al., 2022). Bilabial and labiodental consonants, such as those in /f□fi/ and /vevi/, were more associated with lightness than the alveolar and post-alveolar/velar consonants. Front unrounded vowels were predictive of lightness (as in Walker & Parameswaran, 2019, and Sidhu et al., 2022) as shown in /fIfe/ and /sεse/. Pseudowords rated as heavy were more likely to contain stops and af/fricatives, especially if voiced (consistent with Sidhu et al., 2022; Greenberg & Jenkins, 1966), for example /b/, /g/, and /□/in /bεge/ and /□o□u/. For the place of articulation, the post-alveolar/velar consonants were preferred, over the alveolar and bilabial/labiodental categories, in both consonant positions, for example /g/ and /k/ in /gebi/ and /pIke/. Heaviness was also associated with back rounded vowels, for example /o/, /u/, and /□/, in /□o□u/ and /p□po/, rather than front unrounded vowels (as in Walker & Parameswaran, 2019, and Sidhu et al., 2022).

#### Arousal

For the arousal domain, R^2^ was .48 for the calming scale and .36 for the exciting scale, indicating a reasonable fit between phonetic features and ratings. Sonorants, as in /mulu/ and /lonu/, were strongly associated with calming ratings (Sidhu et al., 2022; see also Aryani et al., 2020) and these examples also illustrate that voiced consonants were more predictive of calming ratings than unvoiced consonants (Aryani et al., 2018). Bilabial/labiodental consonants were also considered calming and, in keeping with the preference for sonorants, six of the ten most calming pseudowords contain the bilabial sonorant /m/ at least once. The back rounded vowels were predictive of calming ratings, as seen in /mumo/ and /m□mo/, and consistent with Sidhu et al. (2022). By contrast, stops and af/fricatives predicted pseudowords rated as exciting (Sidhu et al., 2022; see also Aryani et al., 2020), for example /t/ or /k/ and /z/ or /□/ respectively, in /tIke/ and /z□□i/. Consistent with Aryani et al. (2018), unvoiced, rather than voiced, consonants were associated with exciting ratings. However, there was no significant effect of the place of articulation, with the 10 most exciting pseudowords containing the bilabial /p/, alveolar /t/ and /z/, and postalveolar/velar /□/ and /k/. Front unrounded vowels, as seen in /tIke/ and /z□□i/, also predicted perception of a pseudoword as sounding exciting (Sidhu et al., 2022).

#### Brightness

The brightness domain also exhibited a reasonable fit between phonetic features and ratings, with R^2^ of .47 for the bright scale and .31 for the dark scale. For pseudowords rated as sounding bright, the manner of articulation for consonants was less important than their voicing, consistent with Newman (1933). The unvoiced stops /t/ and /k/, and unvoiced af/fricatives /f/, /s/, and /□/, occurred in nine of the ten brightest pseudowords, while the voiced /v/ and /z/ occurred only once and in the same item: /vizi/. For place of articulation, there were small but significant effects for alveolar and post-alveolar/velar consonants in the first and second consonant positions respectively, for example, /s/ and /□/ in /si□i/. Bright ratings were predicted by the front unrounded vowels (Newman, 1933), for example /e/ and /i/, in /kike/ and /pite/ (note that /vizi/, although an outlier in containing voiced consonants, also contains two front unrounded vowels). For darkness ratings and consonants, however, both manner of articulation and voicing were important. Voiced, rather than unvoiced, consonants were more predictive of darkness ratings (Newman, 1933) and are the only consonants to appear in the ten darkest words. Af/fricatives were associated with dark ratings, but there was only one occurrence in the ten darkest pseudowords – /vu□o/ – although there were a further eight in the next ten items. Back rounded vowels were more predictive of darkness ratings (Newman, 1933), for example in /gubo/. Vowel height had no influence on ratings of either brightness or darkness.

#### Valence

The valence domain also had a reasonable fit between phonetic features and ratings, with R^2^ of .4 for the good scale and .28 for the bad scale. Pseudowords rated as good were associated with sonorants rather than obstruents, as suggested by Adelman et al. (2018), for example, in /meni/ and /lunu/. Additionally, there was a small but significant influence of unvoiced consonants, although there were only 3 occurrences in the first 30 items; this finding was not consistent with Aryani et al. (2018), who found that unvoiced consonants were rated as negatively valenced. The place of articulation was not a significant predictor, with bilabial /m/ and /b/, and alveolar /l/ and /n/, all appearing in the ten most highly-rated good pseudowords. Front unrounded vowels were more predictive of goodness ratings than back rounded vowels (Körner & Rummer, 2022), as exemplified by /l□me/ and /bIbi/. By contrast, stops and af/fricatives indicated badness (Adelman et al., 2018; Uno et al., 2020), for example, /d/ and /□/ in /dudo/ and /fo□u/. The importance of fricatives for badness ratings was in line with Aryani et al. (2018) who reported that hissing sibilants (of which the fricatives /s/ and /z/ are examples) were associated with negative valence. There was no significant influence of voicing: /dudo/and /fo□u/ contain voiced stops and unvoiced af/fricatives respectively. As with the goodness ratings above, this was inconsistent with Aryani et al. (2018) who reported that positive and negative valence were reflected in voiced and unvoiced consonants, respectively. In terms of the place of articulation, there were no significant effects other than for bilabial/labiodentals as the initial consonant, for example /f/, /p/, and /v/ in /fo□u/, /pupo/, and /v□vu/; these examples also illustrate the significant influence of back rounded vowels on the perception of pseudowords as sounding bad (Körner & Rummer, 2022).

#### Size

The size domain had the worst model fit, with R^2^ of .12 for the small scale and .29 for the big scale. Nonetheless, there were some significant phonetic predictors. In contrast to Sidhu et al. (2022), obstruents were preferred over sonorants for smallness ratings but the effect was only significant for (unvoiced) stops, for example /p/ and /t/ in /pete/ and /pIte/ (consistent with Winter & Perlman, 2021), not for af/fricatives. This latter point contrasts with Preziosi & Coane (2017), who reported that fricatives were associated with small size. For the place of articulation, the bilabial/labiodental consonants were more associated with smallness ratings than the alveolar and post-alveolar/velar consonants, for example, /p/ and /f/ in /pete/ and /fIse/. Vowels were not a significant predictor of smallness, again in contrast to previous studies (Sapir, 1929; Auracher, 2017; Sidhu et al., 2022), where smallness was predicted by front vowels. Obstruents were again preferred over sonorants for pseudowords rated as sounding big, but here both (voiced) stops and af/fricatives were significant predictors (as in Sidhu et al., 2022, and consistent with Preziosi & Coane, 2017), for example, /□/ and /b/ in /□I□e/ and /b□ge/. In terms of the place of articulation, the post-alveolar/velar consonants were significant predictors of bigness ratings and in either consonant position: /□/, for example appears in both positions in /□I□e/. Consistent with prior work (Sapir, 1929; Auracher, 2017; Sidhu et al., 2022), big ratings were significantly associated with back rounded vowels, for example /□/ and /o/ in /g□do/, and with mid-height vowels in the second vowel position, as also illustrated by /g□do/.

#### Global observations

Since the same pseudowords were rated across seven different domains, we examined the 10 most highly-rated pseudowords on each of the two scales for each domain (since both scales are used in the phonetic analysis – see below) to see how often pseudowords repeated across domains. It was clear that these highly-rated pseudowords fell into two distinct groups that shared phonetic and acoustic features (see also Lacey et al., 2024). The top ten pseudowords for the rounded, heavy, soft, calming, good, dark, and big scales were largely composed of sonorants, voiced stops and af/fricatives, and back rounded vowels (Table 4A). Acoustically, these pseudowords had a generally steeper spectral tilt, more gradual changes in power, and a smoother, more continuous speech envelope, together with more periodicity and voicing, and less variability (see Lacey et al., 2024 for detailed acoustic analyses). For the pointed, light, hard, exciting, bad, bright, and small scales, however, the most highly-rated pseudowords were largely composed of unvoiced stops and af/fricatives, and front unrounded vowels (Table 4B).

Acoustically, these pseudowords exhibited a generally flatter spectral tilt, more abrupt changes in power, and a discontinuous speech envelope, together with less periodicity and voicing, and more variability (see Lacey et al., 2024). Despite the fact that the pseudowords in each of the two groups of domains were very similar phonetically and acoustically, only eight of the pseudowords repeated within the first group and only seven in the second group (Table 4A & 4B), such that over 70% of the pseudowords in each group were unique.

Finally, as an additional test of domain-general vs domain-specific accounts of sound symbolism, we considered the words used as the endpoints for each domain – i.e., rounded/pointed, light/heavy, hard/soft, and so on. (These appeared with the rating scale on every trial and therefore participants were most frequently exposed to these compared to the additional, explanatory, anchor words that only appeared in the task instructions.) We then extracted the ratings for these endpoint words on scales reflecting arousal (calm/excited), valence (unhappy/happy) and dominance (controlled/in control) from a large-scale rating study of nearly 14,000 words (Warriner et al., 2013). This allowed us to place the endpoints of our domains on continua reflecting two potential domain-general factors and one irrelevant factor (to our knowledge, dominance has not been advanced as an explanation for sound-symbolic associations, and thus serves as a reference condition) according to an independent dataset. We then grouped the endpoints of each domain into high or low arousal, positive or negative valence, and high or low dominance, and counted the incidence of each phonetic feature in the 10 most highly-rated pseudowords at each end of each domain. A domain-general account of sound symbolism would predict that the incidence of each phonetic feature should be similar across domains for each level of the underlying domain-general factor, whether arousal or valence, but should vary across domains for each level of the irrelevant reference factor of dominance. In fact, across domains, we found no coherent pattern of phonetic features that related to arousal, valence, or dominance (Supplementary Tables 5A, B, & C, respectively). Instead, the patterns of phonetic features appeared to be specific to each domain, thus providing evidence against a domain-general account.

## GENERAL DISCUSSION

With few exceptions, studies of sound symbolism have concentrated on a single mapping of sound to meaning, predominantly that of the rounded-pointed dimension of shape. Those studies that have addressed multiple domains of meaning have generally involved a trade-off between either a large number of domains but relatively few real/pseudowords (e.g., Miron, 1961; Johansson et al., 2020; Sidhu et al., 2022) or relatively few domains compared to very large sets of either real words (Winter et al., 2017; see also Tzeng et al., 2017) or pseudowords (Westbury et al., 2018). Here, we obtained ratings across seven domains for a large set of 537 pseudowords as a reasonable compromise; these ratings were used as the outcome variables in regression analyses to determine the phonetic features associated with each domain.

Although the multi-domain studies referred to above were designed to answer particular questions, the approach we took here has some advantages over these earlier studies. Firstly, we collected dimensional ratings of sound-to-meaning correspondence (Miron, 1961; Sidhu et al., 2022) which allow for graded responses across the entire stimulus set, rather than forced choice assessments of meaning (Tzeng et al., 2017) or yes/no category membership decisions (Westbury et al., 2018) which could obscure the granular detail achieved here. Secondly, we extensively sampled the phonological space and ensured that a wide variety of phonetic features were both adequately and equally represented (see McCormick et al., 2015, for details); for example, affricates were not included in the phonetic inventory of Sidhu et al. (2022). The resulting pseudowords were controlled for the number and structure of syllables, for example they contained either voiced or unvoiced consonants but not both (though this might arguably be a limitation given that such instances occur in real words). In common with Johansson et al. (2020) and Westbury et al. (2018), we modelled both manner and place of articulation; the latter was not accounted for by Sidhu et al. (2022). Finally, rating the same pseudowords for multiple domains enables meaningful cross-domain comparisons, as opposed to comparing different domains across different studies that might have different phonetic inventories, stimulus set size, measurements, etc. This approach to both ratings and phonological space allowed us to directly assess the relative contribution of particular phonetic features across multiple domains of meaning.

The ratings analysis revealed many cross-domain associations that could be broadly separated into those reflecting physical, and corresponding perceptual, relationships between object properties, e.g., that between size and weight, and those that were metaphorical in nature, e.g., that between valence and weight (discussed further below). To our knowledge, the present study is the first to directly address the issue of physical/perceptual vs. metaphorical associations in sound-symbolic mappings, discussed in more detail below. Of particular note is the fact that different participants contributed to each of the fourteen scales and were unaware that there was another scale for each domain, or that other domains were being investigated, so to find these relationships suggests a fairly robust phenomenon that merits further research. As discussed below, the phonetic analysis largely replicated, in one multi-domain study, the results of several single-domain studies. Both the ratings and phonetic analyses provided evidence against a domain-general account of sound-to-meaning mapping, suggesting instead that domains need not be associated via ratings and that the patterns of phonetic features were largely domain-specific.

### Ratings: cross-domain comparisons

#### General comments

Among the fully opposing relationships (see boxed results in Table 2), the current findings replicated associations, previously reported in both factor (Miron, 1961; Osgood et al., 1957) and network (Sidhu et al., 2022) analyses, between size and weight, i.e., big/heavy and small/light (Miron, 1961; Sidhu et al., 2022), as well as that between shape and texture (rounded/soft, pointed/hard: Sidhu et al., 2022). However, we did not fully replicate the size-shape association reported by Westbury et al. (2018) in which ‘small’ pseudowords were unconnected to any other category while ‘large’ pseudowords were positively associated with roundness but negatively with sharpness. In the present study, the small scale was indeed uncorrelated with either roundedness or pointedness, but the big scale was positively correlated with both (Table 2). This may reflect the fact that the pseudoword set was sub-optimal for the size domain (see Limitations, below); on the other hand, the size-shape association was not found by Sidhu et al. (2022) either, since these domains loaded onto different factors in that study. Interestingly, in the shape domain for the present study, six of the most highly rated items were replicated from McCormick et al. (2015: Table 4).

#### Physical/perceptual and metaphorical relationships

Although some of the cross-domain relationships have been reported previously, as noted above, they were not further elaborated in terms of physical/perceptual or metaphorical relationships. Generally speaking, we define a cross-domain relationship as physical/perceptual if it is between two of the concrete domains (shape, texture, weight, brightness, or size) and as metaphorical if it is between a concrete domain and an abstract domain (arousal or valence); but this distinction is not always clear-cut. Firstly, arousal can have physical consequences in the context of a concrete domain – for example, the differential effort involved in lifting heavy, compared to light, objects (see the cross-domain calming/light relationship discussed below). Secondly, the format of metaphors is not restricted to a concrete vehicle and an abstract topic (Ritchie, 2013). The interlocking relationships (discussed below) between the brightness domain and the size, weight, and valence domains, illustrate the difficulty; for example, in the dark/heavy metaphor both vehicle and topic relate to concrete domains.

Across domains, there were some ‘double dissociation’ effects, in that ratings on the two scales of particular domain-pairs showed opposing correlations (see boxes in Table 2). For example, for the size and weight domains, ratings on the small scale were significantly positively correlated with those on the light scale, but negatively with those on the heavy scale, while the reverse was true for the big scale, i.e., pseudowords rated as small were also rated as light but not heavy, while pseudowords rated as big were also rated as heavy but not light. The size-weight correlations reflect a physical and perceptual relationship. No meaningful cross-domain relationship was found for size and shape – big ratings were correlated with both rounded and pointed ratings, while small ratings were correlated with neither – likely reflecting the relative independence of physical object size and shape. Other potential physical/perceptual relationships were reflected in pointed pseudowords being rated as hard but not soft, while rounded pseudowords were rated as soft but not hard; and hard pseudowords being rated as heavy but not light, while soft pseudowords were rated as light but not heavy.

Other contrastive correlations reflected metaphorical rather than physical relationships. For texture and valence, for example, ratings on the good scale were positively correlated with softness but negatively with hardness, while the reverse was true for ratings on the bad scale. Similarly, good was also positively correlated with light but negatively with heavy, while bad was positively correlated with heavy but negatively with good. Finally, pointed pseudowords were also rated as bright but not dark, while rounded pseudowords were rated as dark but not bright.

While we adopted a strict criterion for both physical and metaphorical relationships – that both scales for each domain must show significant opposing effects (see boxes in Table 2) – it is also worth looking at relationships where only one scale of each domain shows such effects. While we make some suggestions below as to why these cross-domain relationships did not show fully opposing effects, the simplest explanation is that the study was not designed *a priori* to find these physical and metaphorical relationships. In any event, the relationships discussed below are not invalidated by the lack of a full ‘double dissociation’ effect.

As an example of physical/perceptual relationships, calming ratings were significantly positively correlated with light ratings and significantly negatively correlated with heavy ratings, while exciting ratings did not show any significant relationship to the weight domain. This might reflect the fact that, compared to handling heavy items, light items require less effort, i.e., lower arousal. It is unclear why the exciting scale was unrelated to the weight domain but it may be to do with the fact that the arousal domain – like several others – can be characterized in more than one way. We chose to describe low/high arousal as calming/exciting, respectively, but both these terms are positively valenced. While it makes sense that the low-level arousal involved in handling light items could be viewed positively, it is less clear that the increased effort and arousal involved in handling heavy items would also be viewed this way. If we had characterized arousal as restful/stressful (i.e., positive/negative valence, respectively) we might have seen a double dissociation of the kind discussed above, in which restful ratings were positively correlated with light ratings and negatively with heavy ratings, while stressful ratings were positively correlated with heavy ratings but negatively with light ratings.

Additionally, pointed ratings were significantly positively correlated with bad ratings and significantly negatively correlated with good ratings, while rounded ratings did not show any significant relationship to valence. This might reflect the fact that, compared to rounded items, pointed items have a more obvious capacity to cause pain, i.e., a bad thing and not a good thing. It is unclear why the rounded ratings were unrelated to valence.

For metaphorical instances, there was a single-scale relationship between the brightness and size domains in which dark ratings were significantly positively correlated with ratings on the big scale and significantly negatively with those on the small scale, while bright ratings were not significantly related to either dimension of size (Table 2). This asymmetric relationship is consistent with the fact that big, but not small, objects can block the light and make the environment darker (incidentally, a physical/perceptual relationship). But the dark/big relationship may also be a metaphorical reflection of real-world looming effects in which approaching visual objects increase in apparent size, or ‘loom’ (itself a sound-symbolic word), and are a cue to the threat of imminent collision (Vagnoni et al., 2012). In relation to sound symbolism and language, it is worth noting that looming effects are also elicited in the auditory modality using sounds of increasing intensity (Bach et al., 2009), and are modulated by semantic content: time to collision is perceived as shorter for threatening (snakes, spiders), compared to non-threatening (butterflies, rabbits) items (Vagnoni et al., 2012).

There was also a single-scale relationship between the brightness and weight domains in that dark ratings were significantly positively correlated with heavy ratings and significantly negatively correlated with light ratings, while bright pseudowords were not significantly related to either dimension of weight (Table 2). This might be another aspect of the threat signal posed by large looming objects, i.e. large objects are also likely to be heavy, but it could also reflect the brightness-weight illusion in which dark objects are judged as heavier than light objects (Walker et al., 2010) and the related crossmodal correspondence in which heavier objects are judged as darker (Walker et al., 2017). We should also note that darkness could be metaphorically heavy, in the sense of being oppressive, or in the sense of being so complete as to be almost tangible. For example, speaking about the psychological effects of warfare in his 2009 address to the Veterans of Foreign Wars Convention, former President Barack Obama referred to “the heavy darkness of depression” (Obama, 2009). The word ‘light’ is polysemous, having connotations of brightness as well as weight, which is why we reserved it for one end of the weight domain rather than also using it as one end of the brightness domain (although it was one of the anchor words in the instructions for the bright scale [Supplementary Table 1], it did not appear on the ratings trials themselves). This may be another instance where describing the scale differently might have produced fully opposing effects.

A final example is that goodness was significantly positively correlated with brightness and negatively correlated with darkness. As an example of this metaphorical relationship, we might recall the famous dictum of US Supreme Court Justice (1916-1939) Louis Brandeis who, writing about “publicity [i.e., transparency] as a remedy for social and industrial diseases” remarked that “sunlight is said to be the best disinfectant” (Brandeis, 1913, p10). Additionally, the badness and darkness scales were significantly positively correlated, reflecting a metaphorical equation of darkness with evil as, for example, in references to “the dark side” in the Star Wars universe (e.g., Kurtz & Lucas, 1977). The only element of the double dissociation that is missing is that the bad and bright ratings were negatively correlated, but not significantly so.

There were also cross-domain correlations that could plausibly reflect either physical/perceptual or metaphorical relationships depending on the circumstances. For example, among the contrastive correlations, pseudowords rated as calming were also rated as rounded but not pointed, soft but not hard, and dark but not bright; the reverse was true for pseudowords rated as exciting – these were also rated as pointed but not rounded, hard but not soft, and bright but not dark. Finally, other relationships were harder to interpret; for example, pointed pseudowords were rated as heavy but not light, while big pseudowords were rated as hard but not soft. These do not seem to reflect obvious real-world or perceptual connections, nor to function well as metaphors.

Although they were not the focus of previous studies, several researchers have reported cross-domain associations that reflect physical/perceptual relationships: for example, size-weight (Miron, 1961; Sidhu et al., 2022) as discussed above, but also size-strength (Miron, 1961) in which big/small mapped onto strong/weak, and valence-pleasantness (Miron, 1961; Sidhu et al., 2022) in which good/bad mapped onto pleasant/unpleasant. Previous studies have also reported relationships that we can now characterize as metaphorical or figurative in nature. These include valence-beauty (Sidhu et al., 2022), in which good/bad mapped onto beautiful/ugly, perhaps partly reflecting the figurative expression ‘ugly as sin’. Another example is speed-arousal (Sidhu et al., 2022) in which fast/slow mapped onto to exciting/calming, respectively. The fast-exciting association may reflect risky behavior – ‘fast’ can be defined as “involving or engaging in exciting or shocking activities” (Concise OED, 2011) as in the movie title ‘Fast Times At Ridgemont High’ (Azoff et al., 1982). However, the line between what constitutes a real-world or a metaphorical relationship is sometimes not easily drawn: several studies report a shape-speed relationship in which pointed (sharp)/round map onto fast/slow, respectively (Tzeng et al., 2017; Sidhu et al., 2022). The fast-pointed association could reflect real-world aerodynamic design, but also the metaphorical motion verb ‘to dart’. The gap between physical and metaphorical associations is perhaps bridged by the relationship shown in Sidhu et al. (2022) between tense/relaxed and both inhibited/free (physical) and hard/soft (either physical or metaphorical).

### A grounded account of cross-domain associations

One way in which these cross-domain associations could arise is though grounding in sensorimotor experience (e.g., Barsalou, 2008). This is straightforward for those associations that reflect physical/perceptual relationships, like that between the size and weight domains, but it may also apply to the metaphorical associations. Indeed, Lakoff and Johnson (2003) suggest that knowledge is structured by metaphorical mappings from sensorimotor experience. Take, for example, the cross-domain association in the present study between texture and valence, in which items rated as soft were also rated as good, and those rated hard were also rated bad. This is consistent with a study showing that holding hard objects increased inflexibility in negotiations, and that rough objects made social interactions appear more difficult (Ackerman et al., 2010), suggesting that physical haptic interactions might “scaffold […] conceptual and metaphorical knowledge” (ibid., p1712: note, however, that some of these effects have proved difficult to replicate, see Rabelo et al., 2015). We also observed an association in the present study between weight and valence, in which light was good and heavy was bad. The idea of sin and/or guilt as a burden to be borne is found extensively in the Bible (Lam, 2016); for example, Isaiah 1:4 speaks of being “[…] laden with iniquity […]”, and a later, secular, example occurs in Shakespeare’s *Macbeth*: “The *sin* of my ingratitude even now / was *heavy* on me […]” (Shakespeare (*c*.1606/1984), Macbeth, I_iv_ 15-16, emphasis added).

Similarly, while roundedness had little to do with either of the valence scales, pointedness was strongly associated with ‘bad’ and negatively related to ‘good’. Here, we can note the common expression in English and Korean that morally dubious acts or thoughts ‘prick’ the conscience (Ku et al., 2020). This may also be grounded, as shown by a study in which separate groups of participants were asked to recall instances of their own ethical or unethical actions; to decide between lying or telling the truth; and to make moral judgments on others’ actions. Those who recalled unethical actions were less likely to choose a traditional Korean acupuncture (needle prick) treatment for indigestion, opting instead for a dose of a strong medication. Given an actual needle prick, those who chose to lie reported more pain, and that the needle went deeper, than those who chose to tell the truth and those in a control group; and also rendered stricter moral judgments compared to a control group (Ku et al., 2020). This association is also alluded to in *Macbeth*: “By the *pricking* of my thumbs / Something *wicked* this way comes” (Shakespeare (*c*.1606/1984), Macbeth, IV_i_ 44-45, emphasis added) although it surely has a longer history.

As a final example of grounded metaphorical cross-domain relationships, there was a metaphorical connection between brightness and valence in the present study, in which bright was positively associated with good but negatively with bad, while the reverse was true for dark (see also Meier et al., 2004). That bright is good and dark is bad is a common conceptual metaphor that is not limited to language contexts (for example, see Forceville & Renckens [2013] for the use of visual lightness and darkness in films to indicate valence information about characters and situations). This metaphorical connection may be grounded in experience of the dark as potentially dangerous: in the real world, feelings of personal safety are lower, and the perceived risk and fear of crime are greater, at night than during the day (reviewed in Uttley et al., 2024). A literary example would be “[…] the night is dark and full of terrors […]” (Martin, 1996) and therefore the dark is bad; in this connection, we could also recall Justice Brandeis’s less well-remembered, but metaphorical, remark that “[…] electric light is the best policeman” [Brandeis, 1913]).

Future work should further explore physical vs metaphorical sound-symbolic relationships, not least because some sound-symbolic mappings have yet to be established. Real-world relationships might include loudness and both size (Smith & Sera, 1992) and brightness (Root & Ross, 1965) in which loud/quiet map onto big/small and bright/dark, respectively (see also Spence, 2011). Metaphorical relationships could include hygiene-morality, in which clean/dirty map onto moral/immoral, respectively (see Lee & Schwarz, 2010; Ding et al., 2020); form-honesty, in which straight/crooked are associated with honest/dishonest, respectively (see, for example, Yu, 2016; Zebrowitz et al., 1996; Zhu et al., 2023); and both importance-size and importance-weight, where important/unimportant potentially map onto both big/small and heavy/light, respectively (see Yu et al., 2017). Furthermore, an account of cross-domain associations that is grounded in sensorimotor experience suggests that specific cortical areas might mediate particular sound-symbolic mappings, for example, the lateral occipital complex for shape (reviewed in Lacey & Sathian, 2014) and the parietal operculum for texture (Stilla & Sathian, 2008; Sathian et al., 2011). Such an approach has validated grounded accounts of metaphor comprehension for metaphors related to texture (Lacey et al., 2012), body parts (Lacey et al., 2017), and olfaction (Pomp et al., 2018).

### Phonetic analysis

#### General comments

The present study is not the first to quantify the relative weights of different phonetic features in relation to multiple domains of meaning (see Westbury et al., 2018; Sidhu et al., 2022). However, we modeled both manner and place of articulation (as did Westbury et al., 2018, but not Sidhu et al., 2022), and we did so for a large set of 537 auditory pseudowords. Westbury et al. (2018) presented pseudowords visually – so participants ‘heard’ them only via subvocalization – and individual participants only rated a random subset of 200 items from the full set of 7996. Likewise, although Sidhu et al. (2022) presented auditory pseudowords, participants only rated a random subset of 15 from 40 across all domains. Here, all participants heard and rated the full set of pseudowords. Since we could compare all tested domains across the same set of pseudowords, our results offer a robust comparison with previous studies, and in particular those that only investigated a single domain. As mentioned above, a multi-domain approach avoids the confounds involved in making cross-domain comparisons across separate single-domain studies that may differ in phonetic inventories, stimulus set size, measurements, etc. The contributions of the different phonetic features to the sound-symbolic mapping of the seven domains were largely consistent with previous studies, as summarized below.

Sonorants were associated with roundedness (Köhler, 1929, 1947; Ramachandran & Hubbard, 2011; McCormick et al., 2015; Sidhu et al., 2022), softness (Sidhu et al., 2022; note that this contrasts with Sakamoto & Watanabe, 2018, who found that bilabial stops also indicated softness), lightness (Sidhu et al., 2022), calmness (Sidhu et al., 2022), and goodness (Adelman et al., 2018). Obstruents were associated with pointedness (Köhler, 1929, 1947; Ramachandran & Hubbard, 2011; McCormick et al., 2015; Sidhu et al., 2022), hardness (Sakamoto & Watanabe, 2018; Sidhu et al., 2022), heaviness (Greenberg & Jenkins, 1966), exciting-ness (Aryani et al., 2020; Sidhu et al., 2022), and badness (Adelman et al., 2018; Uno et al., 2020).

Voiced consonants were primarily associated with heaviness (Sidhu et al., 2022), darkness (Newman, 1933), and large size (Westbury et al., 2018) but there were also significant, if weaker, associations with roundedness (Köhler, 1929, 1947; Ramachandran & Hubbard, 2001; McCormick et al., 2015; Sidhu et al., 2022), and calmness (Aryani et al., 2020). Unvoiced consonants were most strongly associated with pointedness (McCormick et al., 2015; Sidhu et al., 2022), lightness (Sidhu et al., 2022), exciting-ness (Aryani et al., 2018), brightness (Newman, 1933) and small size (Winter & Perlman, 2021), with weaker associations to hardness (Sakamoto & Watanabe, 2018), and goodness (see Mooshammer et al., 2023).

For vowels, consistent with previous studies, back/rounded and front/unrounded vowels were respectively associated with roundedness and pointedness (Köhler, 1929, 1947; Ramachandran & Hubbard, 2001; McCormick et al., 2015), heaviness and lightness (Walker & Parameswaran, 2019; Sidhu et al., 2022), calming and exciting (Sidhu et al., 2022), darkness and brightness (Newman, 1933), and badness and goodness (Körner & Rummer, 2022). Back/rounded vowels were also associated with softness, a new finding and in contrast to Sidhu et al. (2022) who found no association between vowels and either hardness or softness. Consistent with earlier findings (Sapir, 1929; Auracher 2017; Sidhu et al., 2022), back/rounded vowels were associated with large size but we did not find the corresponding association between front/unrounded vowels and small size.

In terms of the place of articulation, bilabial/labiodental consonants were associated with roundedness (see also Westbury et al., 2018), softness, lightness, and calmness. Both alveolar and postalveolar/velar consonants were associated with pointedness (see also Westbury et al., 2018), hardness, and brightness, while only post-alveolar/velar consonants were associated with heaviness and also large size. The functional importance of the place of articulation is that it alters the length of the vocal tract involved in producing the sound and thus has acoustic consequences (Reetz & Jongman, 2020). In brief, bilabial/labiodental consonants result in low-frequency sounds while alveolar consonants result in high-frequency sounds, and intermediate frequencies arise from post-alveolar/velar consonants (for a more detailed treatment of the acoustic consequences of the place of articulation, see Lacey et al., 2024). In this respect, it is interesting that roundedness was associated with the low-frequency bilabial/labiodental consonants while pointedness was associated with the relatively higher-frequency alveolar and post-alveolar/velar consonants. These frequency differences match those for vowel sounds associated with rounded/pointed pseudowords (Knoeferle et al., 2017) and visual shapes (Parise & Pavani, 2011). Similarly, the association of brightness with the relatively higher-frequency alveolar and post-alveolar/velar consonants is consistent with a previous finding that pseudowords are produced with a higher fundamental frequency for brighter, compared to darker, colors (Tzeng et al., 2018). Other associations, reported here for the first time, make intuitive sense; for example, that low-frequency bilabial/labiodental consonants sound calming and soft, while the relatively higher-frequency alveolar and post-alveolar/velar consonants sound hard.

For vowel articulation, backness/rounding was an important contributor to almost every domain as described above. Vowel height, however, was relatively unimportant, being non-significant for 18 of the 28 scales (64%), and with small beta coefficients of ≤ .1 (Table 3), although it must be noted that there were no low vowels in the phoneme set, which contained only high and mid-height vowels (Table 1).

#### Phonetic analysis: cross-domain comparisons

Comparing across domains, ratings were best predicted by collections of phonetic features that were relatively specific to each domain. For example, in the shape and weight domains, one scale (rounded or light) was associated with sonorant while the other (pointed or heavy) was associated with obstruents. But, while they were broadly similar in terms of consonants, these scales differed in their vowel associations: back vowels for the rounded and heavy scales, and front vowels for the pointed and light scales (see Table 3). However, the texture and size domains showed more differences: hardness was indicated by obstruents and unvoiced consonants but not vowels, while softness was indicated by sonorants (and so unvoiced consonants were not relevant), and by back rounded vowels. In the size domain, bigness was associated with obstruents, voiced consonants, and back rounded vowels, while smallness was indicated only by stops and unvoiced consonants, not by af/fricatives or vowels. Even positive correlations in ratings across domains were no guarantee of similarity in the related phonetic features. On the one hand, the positively correlated rounded/calming and pointed/exciting scales were reflected in common phonetic features: sonorants, voiced consonants, and back rounded vowels for rounded/calming, and obstruents, unvoiced consonants, and front unrounded vowels for pointed/exciting. By contrast, although the soft/light and hard/heavy scales were both positively correlated and both relied on the same consonants, the ratings combined with vowels differently: back rounded for softness and front unrounded for lightness, while hardness was not reflected in vowels at all and heaviness was associated with back rounded vowels. Finally, we should note that the graded nature of the ratings suggests that, in addition to the individual phonetic features of each pseudoword, participants’ judgments likely involved processing/analysis at the segment or even whole-item level (McCormick et al., 2015; see also Thompson & Estes, 2011).

### Sound symbolism and phonaesthemes

The relationship between the shape and brightness ratings for pseudowords suggests that ‘pointed’ and ‘bright’, and ‘rounded’ and ‘dark’, go together. This seems intuitively correct as reflecting both a physical and a metaphorical relationship, but it is also interesting in relation to real words and, in particular, those characterized by phonaesthemes. Phonaesthemes are an example of systematic non-arbitrary associations to meaning and have not traditionally been assumed to exhibit iconicity; for example, /tw/ in ‘tweak’, ‘twiddle’, ‘twine’, twirl’, and ‘twist’ is associated with a twisting/turning movement (see Nuckolls, 1995), while /sn/ occurs in many words related to the nose or mouth, for example, ‘snout’, ‘sniff’, ‘sneeze’, and ‘snack’, ‘snarl’ (see Bergen, 2004). These phonaesthemic non-arbitrary sound-meaning correspondences may be worth exploring here because, although they arguably arise from a different – probably etymological – mechanism, they are often combined with sound-symbolic segments and, like sound symbolism, they occur across a wide range of languages (see Bergen, 2004).

In terms of a real-life shape-brightness relationship, ‘glint’, ‘glisten’, and ‘glitter’ all refer to actual light conditions where the light is concentrated: ‘small flashes of light’ or ‘sparkling’ light (Concise OED, 2011). These real words also contain the unvoiced consonants and stops (/s/, /t/) and the unrounded vowel /I/ associated with pointedness in our pseudowords. Conversely, words like ‘glimmer’, ‘glow’, and ‘gloom’ refer to conditions in which the light is more diffuse; these real words contain the sonorants (/m/) and rounded vowels (diphthongized /o/ and /□/ in ‘glow’, /u/) that were associated with roundedness in our pseudowords. All these words share the initial phonaestheme /gl/, which is found in a number of words whose meaning relates to light, as above, or vision, for example ‘glimpse’ and ‘glance’ (Bergen, 2004), and which is potentially traceable to the Indo-European prefix ‘*ghlei-*’ meaning ‘shine; glitter; glow’ (Klein, 1966). Interestingly, the same phonaestheme begins words that are positively or negatively valenced but metaphorically bright or dark, such as ‘glad’, ‘glee’, or ‘glum’, and that also contain phonemes associated with brightness and darkness (see the discussion of the brightness-valence metaphor above). ‘Glad’ derives from Old English ‘gl d’ meaning ‘bright’ or ‘shining’ (Concise OED, 2011) and perhaps ultimately from another Indo-European root, the prefix ‘*ghledho-*’, also meaning ‘bright’ (Klein, 1966), while ‘glee’ is ultimately related to the Old Norse ‘*g□*ā’ meaning ‘to shine’ (Klein, 1966). Both ‘glad’ and ‘glee’ contain front unrounded vowels (/æ/ and /i/, respectively) associated with both brightness and positive valence (Table 3), although /æ/ was not included in the vowels tested here. The vowel /□/ in ‘glum’ was also not included in the current study but is sound symbolically associated with negative emotional valence (Yu et al., 2021) and, as another back mid-height vowel, is arguably also a ‘dark’ phoneme (see Newman, 1933).

Some of the sounds that occur in words to do with light also occur in words related to uncleanliness, but are differentiated by appending a different phonaestheme; for example, /gr/ in greasy, grimy, gritty, grotty, grody, and grubby. These suffixes are potentially sound-symbolic, although the clean/dirty domain does not seem to have received much attention. In relation to Japanese mimetic words, Miyake (2008) suggests that unvoiced and voiced consonants map onto clean/dirty, respectively; but this is only partially borne out in the examples given above. Nonetheless, the sound-symbolic nature of the suffixes associated with a common phonaesthemic prefix for words related to a common domain of meaning may be worth investigating^1^. Along these lines, while the exact role of phonaesthemes in linguistics is still unclear (see Bergen, 2004; Svantesson, 2017; Philps, 2023), they may be a form of organizing principle (Bergen, 2004) in that shared meaning contributes to shared form and vice versa. Uniting phonaesthemes with sound symbolism would be one way in which this could be achieved, i.e., by adding a sound-symbolic suffix to a meaning-specific root, as in /gl/oom or /gl/isten (note that phonaesthemes are involved in the creation of neologisms, see Bergen [2004]; Blust [2003]). Thus, within a group linked by the same phonaestheme, we see iconicity emerging to nuance, for example, the quality of light. Recognizing that phonaesthemes extend to metaphorical usages of the underlying domain of meaning would have two effects. Firstly, it would resolve doubts about group membership; for example, Joseph (2013) comments that it is unclear why ‘glad’ should belong to the /gl/-light group. Secondly, it would enlarge group membership – for example, ‘glad’, glee’, and ‘glum’, were not included in the list of /gl/-light words in Bergen (2004: Appendix A) – and by thus excluding metaphorically related items, the frequency of phonaesthemes in the lexicon, and the connection between sound symbolism and phonaesthemes as an organizing principle, may be underestimated.

### Cross-domain associations and the origins of language

The cross-domain ratings relationships between dark and big, and between bad and both dark and pointed (discussed above, and see Table 2) are interesting in view of the long-standing conjecture that sound symbolism is a candidate for the emergence of referential meaning in the origins of human language and vocal communication (see, for example, Swadesh, 1971; Mithen, 2005; Perniss & Vigliocco, 2014). The threat of danger would be among the more important messages to communicate and indicating looming threats via the dark-big-bad association might achieve this. In the same vein, pointed pseudowords were also rated as bad but not good, while rounded pseudowords were unrelated to either dimension of valence, and pointedness would most often indicate a threat (for example, claws, fangs, talons, teeth, thorns, spines, etc.). In terms of how vocabulary might have evolved, Table 4 suggests some degree of specificity, given that roughly three-quarters of the most highly-rated pseudowords were unique, regardless of whether the underlying ratings were correlated or not. For example, ratings on the dark, big, and bad scales were positively correlated with each other; but while the most highly-rated pseudowords on each of these scales generally have back rounded vowels in common, for the most part, dark is differentiated from big by voiced stops for the former and affricates (/□/, /□/) for the latter, and from bad by unvoiced stops and fricatives (/f/, /v/, /s/, /z/). Similarly, although pointed and bad ratings were also correlated and the pseudowords have unvoiced stops in common, they are differentiated by front unrounded vowels for the highly-rated pointed pseudowords and back rounded vowels for the highly-rated bad pseudowords. By contrast, ratings on the rounded and bad scales were uncorrelated but the most highly-rated rounded pseudowords were mainly characterized by voiced stops and sonorants, whereas the most highly-rated bad pseudowords were associated with unvoiced stops and af/fricatives; back rounded vowels were common to both scales, though much more important to the round scale. One implication of this is that, during the evolution of language, sound-to-meaning iconicity may have been co-opted in a domain-specific manner (see Lacey et al., 2024, for a discussion of how this might have been enabled by evolutionary changes in the larynx).

### Domain-general vs domain-specific accounts of sound symbolism

Both the ratings and the phonetic analyses argue against domain-general accounts of sound symbolism based on, for example, mappings to higher-order or abstract dimensions such as arousal (Aryani et al., 2020) or valence (for example, Osgood, 1969; but see Tzeng et al., 2017, for commentary). The claim that sound and meaning are connected via arousal (Aryani et al., 2020) is undermined by the fact that only the shape domain was tested when, if true, the effect should generalize to other domains. The present results allow a test of this because a strong prediction of this account would be that arousal ratings for the pseudowords would be related to their ratings for any other domain. In other words, the opposing ends of every domain would be assigned to either low or high arousal – for example, dark and bright, respectively – and we would expect that low/high arousal ratings would be positively correlated with one and negatively with the other. For brightness, shape and texture, this was, in fact, the case (Table 2) but, it was not so for the other domains. Thus, the results suggest that arousal is unlikely to be a domain-general factor underlying all sound-symbolic mappings. This was also the case for valence ratings: these showed fully opposing correlations for only two of the seven domains (Table 2). The lack of evidence for a domain-general account based on the ratings data is consistent with Tzeng et al. (2017) who analyzed their ratings data for semantic relatedness to a range of possible higher-order factors but found no convincing evidence for a domain-general factor underlying the sound-symbolic mappings tested.

Domain-general accounts are also contradicted by the finding that different domains were underpinned by different patterns of phonetic features (Table 3). Even where there were cross-domain correlations for ratings, the phonetic patterns were not necessarily related: some correlated domains shared similar phonetic features (sometimes very similar), but many did not. For an arousal account in particular, ratings for exciting (high arousal) pseudowords were significantly associated with obstruents which was also true for ratings on other high-arousal scales – for example, pointed, hard, and heavy – but not true for the high-arousal bright scale. In the same way, exciting ratings were significantly associated with unvoiced consonants, as were pointed and bright ratings, but ratings for the high-arousal dimension of heaviness were associated with voiced consonants. In addition, we grouped the 10 most highly-rated pseudowords by reference to the high/low arousal and positive/negative valence ends of each domain, but found no coherent pattern of phonetic features that would suggest a domain-general account of sound symbolism (Supplementary Tables 5A & B). For further evidence against a domain-general account in terms of the acoustic parameters associated with different domains, so far untested in this respect, see Lacey et al. (2024).

### Limitations

A general caveat is that the pseudowords were constructed using phonemes that had established associations to the rounded-pointed dimension of shape (McCormick et al., 2015) and thus might be said to be optimized for the shape domain in preference to the others tested here. Against that, as the survey of prior literature shows (see Introduction), none of these phonemes are *exclusively* associated with shape but, instead, have connections to several other domains. Nonetheless, the pseudowords only employ a subset of all English phonemes. In particular, /a/, /[r]/ and / /, which have established associations to the size domain (reviewed in Ekström, 2022), are absent, which may help explain the poor model fit for size (Table 3). In other words, it may be that the pseudowords are not so much optimal for shape but rather sub-optimal for size. In fact, Ekström (2022) suggests that there is no settled account of size sound symbolism. In any case, a more complete account of the phonetic bases for different sound-symbolic domains may emerge from either more targeted sets of phonemes or, given that phonemic associations are likely not exclusive, from a pseudoword set that samples the complete phonetic space with each phoneme equally represented (or in proportion to their frequency in the relevant language).

A conceivable limitation for the shape domain is that the speaker who recorded the pseudowords might have unconsciously pronounced them differently according to their expectations about the pseudowords’ roundedness/pointedness (see also Lacey et al., 2020, 2024). This is unlikely, however, because the speaker was instructed to employ a neutral intonation and two independent judges rejected items that did not sound both neutral and consistent with the other recordings. For the other domains reported here, the speaker was unaware that their recordings would be used for other domains in the future, therefore they cannot have been influenced by prior knowledge or expectations about those domains.

In terms of the trade-off between the number of domains and pseudowords, studies using a small pseudoword set may be at a disadvantage: the smaller the pseudoword set, the more items repeat across domains because participants have a restricted choice. For the set of 537 pseudowords used here, we observed more than 70% unique items in the 10 most highly rated items for both scales in 7 domains. For a smaller set of 40 pseudowords, Sidhu et al. (2022, Figure 4) provide information for 4 of the 24 domains: for the 8 most highly rated items at the opposing ends of each domain, less than 25% are unique (and even in the 4 most highly rated items, unique items still constitute less than 50%). By contrast, for a large set of 7996 pseudowords reported in Westbury et al., (2018: Tables 4, 7, 9), the 10 items most likely to be assigned to 3 domains were 100% unique. (There may be many reasons for this last result aside from the very large set size: for example, pseudowords could be more than two syllables and were not restricted to a CVCV sequence since they could start with a vowel and end with a consonant, and could also include biphones like /br/ and /mb/.)

## CONCLUSIONS

The present findings suggest that the contributions and relative weightings of different phonetic features are largely specific to each domain and thus argue against a domain-general account in which sound-symbolic mappings are driven by a single common factor. For a similar conclusion from an acoustic perspective, see Lacey et al. (2024). The results also suggest that many sound-symbolic mappings may be related to each other in ways that reflect relationships that are either physical and perceptual or metaphorical and figurative. This might be a more parsimonious account than grouping domains together under over-arching factors such as activity or potency – which are essentially themselves ‘super’-domains – and would reflect two basic modes of expression: literal and figurative language. Further work is needed to explore this more fully.

## ACKNOWLEDGMENTS

This work was supported by grants to KS and LCN from the National Eye Institute at the NIH (R01EY025978) and the Emory University Research Council. An earlier version of these data was presented at the 2022 meeting of the Cognitive Neuroscience Society (Lacey et al., 2022) and was available as a preprint at *bioRxiv*, doi: [TBA]. We thank Nancy Bliwise, Department of Psychology, Emory University, for statistical advice.

Author contributions: SL, KS and LCN designed the study; KLM collected the data; KLM and SL analyzed the data; and SL, KLM, KS and LCN wrote the paper.

## DATA AVAILABILITY

The set of 537 pseudowords is available at https://osf.io/ekpgh/ and the rating data for all domains are available at https://osf.io/y9zjc/.

## Footnotes

1 We should note that phonaesthemes can be suffixes as well as prefixes (Otis & Sagi, 2008), for example, ‘bump’, ‘dump’, ‘lump’, and ‘thump’.

