## Supplementary Material for "Phonetic underpinnings of sound symbolism across multiple domains of meaning"

PHONETIC UNDERPINNINGS OF CROSS-DOMAIN ASSOCIATIONS

IN SOUND SYMBOLISM

| Lynne C. Nygaard  Department of Psychology  Emory University  College of Arts and Sciences  Atlanta, GA 30322, USA  Tel: 404-727-0766  Fax: 717-531-0384  | K. Sathian  Department of Neurology  Milton S. Hershey Medical Center  Penn State College of Medicine  Hershey, PA 17033-0859, USA  Tel: 717-531-1801  Fax: 717-531-0384  |
| --- | --- |

**Supplementary Table 1:** Anchor words for the categorical opposites for each scale.

| **Domain** | **Categorical opposites** | **Anchor words** |
| --- | --- | --- |
| Shape | Rounded | rounded; bloblike; amoeboid |
|  | Pointed | pointed; angular; jagged |
| Texture | Hard | hard; unyielding; firm |
|  | Soft | soft; squishy; cushioned |
| Weight | Light | light; weightless; airy |
|  | Heavy | heavy; weighty; hefty |
| Size | Small | small; tiny; little |
|  | Big | big; large; huge |
| Brightness | Bright | bright; light; luminous |
|  | Dark | dark; dim; unlit |
| Arousal | Calming | calming; tranquil; passive |
|  | Exciting | exciting; exhilarating; active |
| Valence | Good | good; nice; excellent |
|  | Bad | bad; nasty; terrible |

**Supplementary Table 2:** Multicollinearity statistics; VIF = variance inflation factor.

| **Predictor** | **Tolerance** | **VIF** |
| --- | --- | --- |
| Stops | .39 | 2.54 |
| Af/fricatives | .39 | 2.55 |
| Unvoiced | .84 | 1.19 |
| Alveolar 1 | .72 | 1.39 |
| Alveolar 2 | .74 | 1.35 |
| Post-alveolar/velar 1 | .71 | 1.41 |
| Post-alveolar/velar 2 | .72 | 1.38 |
| Back rounded | .95 | 1.05 |
| High 1 | .94 | 1.06 |
| High 2 | .98 | 1.02 |

**Supplementary Table 3:** Durbin-Watson test values.

| **Domain** | **Scale** | **Test value** |
| --- | --- | --- |
| Shape | Rounded | 2.23 |
|  | Pointed | 2.17 |
| Texture | Hard | 1.89 |
|  | Soft | 1.84 |
| Weight | Light | 1.94 |
|  | Heavy | 1.81 |
| Arousal | Calming | 1.79 |
|  | Exciting | 1.9 |
| Brightness | Bright | 2.1 |
|  | Dark | 2.01 |
| Valence | Good | 1.93 |
|  | Bad | 2.02 |
| Size | Small | 1.89 |
|  | Big | 1.89 |

**Supplementary Table 4**: B, unstandardized beta value; SE, standard error; β, standardized beta coefficients; 1,2, first or second consonant or vowel position; * p < .05, ** p < .01, *** p < .001.

| **Shape** | Rounded (R^2^ = .81) | | | Pointed (R^2^ = .76) | | |
| --- | --- | --- | --- | --- | --- | --- |
|  | B | SE B | β | B | SE B | β |
| Constant | 3.56 | .05 |  | 2.58 | .05 |  |
| Stops | -0.27 | .05 | -.18*** | 0.94 | .05 | .69*** |
| Af/fricatives | -0.35 | .05 | -.23*** | 0.67 | .05 | .50*** |
| Unvoiced | -0.11 | .03 | -.07*** | 0.52 | .03 | .38*** |
| Alveolar 1 | -0.20 | .03 | -.13*** | 0.12 | .04 | .09*** |
| Alveolar 2 | -0.17 | .03 | -.11*** | 0.18 | .03 | .13*** |
| Post-alveolar/velar 1 | -0.06 | .04 | -.04 | 0.29 | .04 | .20*** |
| Post-alveolar/velar 2 | -0.24 | .04 | -.14*** | 0.41 | .04 | .28*** |
| Back rounded | 1.28 | .03 | .87*** | -0.16 | .03 | -.12*** |
| High 1 | -0.08 | .03 | -.05** | 0.07 | .03 | .05* |
| High 2 | -0.14 | .03 | -.09*** | < -0.01 | .03 | < .001 |

| **Texture** | Hard (R^2^ = .68) | | | Soft (R^2^ = .33) | | |
| --- | --- | --- | --- | --- | --- | --- |
|  | B | SE B | β | B | SE B | β |
| Constant | 3.16 | .04 |  | 4.32 | .05 |  |
| Stops | 0.76 | .04 | .81*** | -0.43 | .05 | -.55*** |
| Af/fricatives | 0.34 | .04 | .36*** | -0.18 | .04 | -.24*** |
| Unvoiced | 0.06 | .03 | .06* | 0.01 | .03 | .01 |
| Alveolar 1 | 0.04 | .03 | .04 | -0.12 | .03 | -.16*** |
| Alveolar 2 | 0.12 | .03 | .13*** | -0.17 | .03 | -.21*** |
| Post-alveolar/velar 1 | 0.21 | .03 | .20*** | -0.20 | .04 | -.23*** |
| Post-alveolar/velar 2 | 0.37 | .03 | .36*** | -0.21 | .04 | -.25*** |
| Back rounded | -0.04 | .02 | -.05 | 0.14 | .03 | .19*** |
| High 1 | -0.03 | .02 | -.03 | -0.06 | .03 | -.08* |
| High 2 | -0.06 | .02 | -.07** | -0.01 | .03 | -.01 |

| **Weight** | Light (R^2^ = .43) | | | Heavy (R^2^ = .53) | | |
| --- | --- | --- | --- | --- | --- | --- |
|  | B | SE B | β | B | SE B | β |
| Constant | 4.25 | .04 |  | 3.55 | .05 |  |
| Stops | -0.52 | .04 | -.74*** | 0.71 | .04 | .82*** |
| Af/fricatives | -0.35 | .04 | -.50*** | 0.38 | .04 | .45*** |
| Unvoiced | 0.11 | .03 | .15*** | -0.22 | .03 | -.25*** |
| Alveolar 1 | -0.09 | .03 | -.13*** | -0.03 | .03 | -.04 |
| Alveolar 2 | -0.11 | .03 | -.15*** | 0.03 | .03 | .03 |
| Post-alveolar/velar 1 | -0.14 | .03 | -.19*** | 0.12 | .03 | .13*** |
| Post-alveolar/velar 2 | -0.22 | .03 | -.29*** | 0.22 | .03 | .23*** |
| Back rounded | -0.1 | .02 | -.14*** | 0.10 | .03 | .12*** |
| High 1 | -0.02 | .02 | -.03 | 0.02 | .03 | .02 |
| High 2 | 0.06 | .02 | .09* | -0.08 | .03 | -.09** |

**Supplementary Table [x] cont.**

| **Arousal** | Calming (R^2^ = .48) | | | Exciting (R^2^ = .36) | | |
| --- | --- | --- | --- | --- | --- | --- |
|  | B | SE B | β | B | SE B | β |
| Constant | 4.51 | .04 |  | 3.52 | .05 |  |
| Stops | -0.47 | .04 | -.62*** | 0.13 | .04 | .18** |
| Af/fricatives | -0.41 | .04 | -.55*** | 0.32 | .04 | .43*** |
| Unvoiced | -0.07 | .03 | -.09** | 0.1 | .03 | .13** |
| Alveolar 1 | -0.04 | .03 | -.06 | 0.04 | .03 | .06 |
| Alveolar 2 | -0.08 | .03 | -.11** | 0.04 | .03 | .06 |
| Post-alveolar/velar 1 | -0.11 | .03 | -.14*** | 0.01 | .03 | .02 |
| Post-alveolar/velar 2 | -0.23 | .03 | -.28*** | 0.04 | .03 | .05 |
| Back rounded | 0.17 | .02 | .22*** | -0.33 | .03 | -.45*** |
| High 1 | < -0.01 | .02 | -.01 | 0.09 | .03 | .11** |
| High 2 | 0.04 | .02 | .05 | < 0.01 | .03 | < .01 |

| **Brightness** | Bright (R^2^ = .47) | | | Dark (R^2^ = .31) | | |
| --- | --- | --- | --- | --- | --- | --- |
|  | B | SE B | β | B | SE B | β |
| Constant | 4.03 | .05 |  | 3.43 | .05 |  |
| Stops | 0.05 | .05 | .05 | 0.09 | .05 | .11 |
| Af/fricatives | 0.01 | .05 | .01 | 0.12 | .04 | .15** |
| Unvoiced | 0.22 | .03 | .25*** | -0.24 | .03 | -.31*** |
| Alveolar 1 | 0.03 | .03 | .03 | < 0.01 | .03 | -.01 |
| Alveolar 2 | 0.07 | .03 | .08* | < 0.01 | .03 | -.01 |
| Post-alveolar/velar 1 | 0.1 | .04 | .1** | -0.06 | .04 | -.07 |
| Post-alveolar/velar 2 | 0.07 | .04 | .07 | 0.01 | .04 | -.01 |
| Back rounded | -0.56 | .03 | -.63*** | 0.36 | .03 | .47*** |
| High 1 | 0.03 | .03 | .04 | -0.03 | .03 | -.04 |
| High 2 | 0.04 | .03 | .04 | 0.04 | .03 | .05 |

| **Valence** | Good (R^2^ = .4) | | | Bad (R^2^ = .28) | | |
| --- | --- | --- | --- | --- | --- | --- |
|  | B | SE B | β | B | SE B | β |
| Constant | 4.73 | .05 |  | 3.01 | .04 |  |
| Stops | -0.52 | .04 | -.66*** | 0.31 | .04 | .5*** |
| Af/fricatives | -0.57 | .04 | -.74*** | 0.32 | .04 | .52*** |
| Unvoiced | 0.08 | .03 | .1** | 0.03 | .03 | .04 |
| Alveolar 1 | 0.03 | .03 | .04 | -0.12 | .03 | -.19*** |
| Alveolar 2 | -0.01 | .03 | -.02 | < -0.01 | .03 | -.01 |
| Post-alveolar/velar 1 | -0.02 | .03 | -.02 | -0.05 | .03 | -.08 |
| Post-alveolar/velar 2 | -0.05 | .03 | -.05 | 0.01 | .03 | .02 |
| Back rounded | -0.21 | .03 | -.27*** | 0.12 | .02 | .19*** |
| High 1 | 0.01 | .03 | .01 | -0.01 | .02 | -.02 |
| High 2 | -0.07 | .03 | -.1** | 0.02 | .02 | .03 |

| **Size** | Small (R^2^ = .12) | | | Big (R^2^ = .29) | | |
| --- | --- | --- | --- | --- | --- | --- |
|  | B | SE B | β | B | SE B | β |
| Constant | 3.43 | .05 |  | 3.66 | .04 |  |
| Stops | 0.11 | .04 | .17** | 0.27 | .04 | .41*** |
| Af/fricatives | -0.03 | .04 | -.05 | 0.29 | .04 | .43*** |
| Unvoiced | 0.12 | .03 | .18*** | -0.13 | .03 | -.2*** |
| Alveolar 1 | -0.03 | .03 | -.05 | 0.03 | .03 | .05 |
| Alveolar 2 | -0.05 | .03 | -.08 | < 0.01 | .03 | .01 |
| Post-alveolar/velar 1 | -0.17 | .04 | -.23*** | 0.15 | .03 | .20*** |
| Post-alveolar/velar 2 | -0.1 | .04 | -.13** | 0.11 | .03 | .15** |
| Back rounded | -0.04 | .03 | -.06 | 0.18 | .03 | .28*** |
| High 1 | -0.03 | .03 | -.05 | -0.02 | .03 | -.02 |
| High 2 | 0.04 | .03 | .05 | -0.09 | .02 | -.14*** |

**Supplementary Table 5:** Percentage count of the phonetic features of the 10 most highly-rated pseudowords using the single, recoded, scale for each domain, grouped by potential domain-general factors, either (A) arousal or (B) valence, and by an irrelevant control factor (C) dominance; this shows that there is no common pattern of phonetic features across domains for any grouping.

| **(A) Arousal** | |  | |  | **Shape** | | **Weight** | | **Texture** | | **Arousal** | | **Valence** | | **Brightness** | | **Size** | |
| --- | --- | --- | --- | --- | --- | --- | --- | --- | --- | --- | --- | --- | --- | --- | --- | --- | --- | --- |
|  | |  | | Low arousal | Rounded | | Light | | Soft | | Calming | | Good | | Dark | | Small | |
| **Consonants (%)** | |  | |  |  | |  | |  | |  | |  | |  | |  | |
| Sonorants | |  | |  | 100 | | 70 | | 60 | | 100 | | 90 | | 30 | | 10 | |
| Stops | | Voiced | |  |  | |  | |  | |  | | 10 | | 70 | | 10 | |
|  | | Unvoiced | |  |  | |  | |  | |  | |  | |  | | 40 | |
| Af/fricatives | | Voiced | |  |  | |  | |  | |  | |  | |  | | 30 | |
|  | | Unvoiced | |  |  | | 30 | | 40 | |  | |  | |  | | 10 | |
| **Vowels (%)** | |  | |  |  | |  | |  | |  | |  | |  | |  | |
| Back rounded | |  | |  | 100 | | 20 | | 50 | | 100 | | 10 | | 80 | |  | |
| Front unrounded | |  | |  |  | | 80 | | 50 | |  | | 90 | | 20 | | 100 | |
|  | |  | | High arousal | Pointed | | Heavy | | Hard | | Exciting | | Bad | | Bright | | Big | |
| **Consonants (%)** | |  | |  |  | |  | |  | |  | |  | |  | |  | |
| Sonorants | |  | |  |  | |  | |  | |  | |  | |  | |  | |
| Stops | | Voiced | |  |  | | 60 | | 20 | | 10 | | 10 | |  | | 20 | |
|  | | Unvoiced | |  | 90 | | 30 | | 80 | | 50 | | 40 | | 80 | |  | |
| Af/fricatives | | Voiced | |  |  | | 10 | |  | | 30 | | 30 | | 10 | | 60 | |
|  | | Unvoiced | |  | 10 | |  | |  | | 10 | | 20 | | 10 | | 20 | |
| **Vowels (%)** | |  | |  |  | |  | |  | |  | |  | |  | |  | |
| Back rounded | |  | |  |  | | 50 | | 30 | | 10 | | 90 | |  | | 30 | |
| Front unrounded | |  | |  | 100 | | 50 | | 70 | | 90 | | 10 | | 100 | | 70 | |
| **(B) Valence** |  | |  | | | **Shape** | | **Weight** | | **Texture** | | **Arousal** | | **Valence** | | **Brightness** | | **Size** |
|  |  | | Negative valence | | | Pointed | | Heavy | | Hard | | Calming | | Bad | | Dark | | Big |
| **Consonants (%)** |  | |  | | |  | |  | |  | |  | |  | |  | |  |
| Sonorants |  | |  | | |  | |  | |  | | 100 | |  | | 30 | |  |
| Stops | Voiced | |  | | |  | | 60 | | 20 | |  | | 10 | | 70 | | 20 |
|  | Unvoiced | |  | | | 90 | | 30 | | 80 | |  | | 40 | |  | |  |
| Af/fricatives | Voiced | |  | | |  | | 10 | |  | |  | | 30 | |  | | 60 |
|  | Unvoiced | |  | | | 10 | |  | |  | |  | | 20 | |  | | 20 |
| **Vowels (%)** |  | |  | | |  | |  | |  | |  | |  | |  | |  |
| Back rounded |  | |  | | | 0 | | 50 | | 30 | | 100 | | 90 | | 80 | | 30 |
| Front unrounded |  | |  | | | 100 | | 50 | | 70 | | 0 | | 10 | | 20 | | 70 |
|  |  | | Positive valence | | | Rounded | | Light | | Soft | | Exciting | | Good | | Bright | | Small |
| **Consonants (%)** |  | |  | | |  | |  | |  | |  | |  | |  | |  |
| Sonorants |  | |  | | | 100 | | 70 | | 60 | |  | | 90 | |  | |  |
| Stops | Voiced | |  | | |  | |  | |  | | 10 | | 10 | |  | | 20 |
|  | Unvoiced | |  | | |  | |  | |  | | 50 | |  | | 80 | |  |
| Af/fricatives | Voiced | |  | | |  | |  | |  | | 30 | |  | | 10 | | 60 |
|  | Unvoiced | |  | | |  | | 30 | | 40 | | 10 | |  | | 10 | | 20 |
| **Vowels (%)** |  | |  | | |  | |  | |  | |  | |  | |  | |  |
| Back rounded |  | |  | | | 100 | | 20 | | 50 | | 10 | | 10 | |  | | 30 |
| Front unrounded |  | |  | | |  | | 80 | | 50 | | 90 | | 90 | | 100 | | 70 |

| **(C) Dominance** |  |  | **Shape** | **Weight** | **Texture** | **Arousal** | **Valence** | **Brightness** | **Size** |
| --- | --- | --- | --- | --- | --- | --- | --- | --- | --- |
|  |  | Low dominance | Rounded | Heavy | Hard | Exciting | Bad | Dark | Small |
| **Consonants (%)** |  |  |  |  |  |  |  |  |  |
| Sonorants |  |  | 100 |  |  |  |  | 30 |  |
| Stops | Voiced |  |  | 60 | 20 | 10 | 10 | 70 | 20 |
|  | Unvoiced |  |  | 30 | 80 | 50 | 40 |  |  |
| Af/fricatives | Voiced |  |  | 10 |  | 30 | 30 |  | 60 |
|  | Unvoiced |  |  |  |  | 10 | 20 |  | 20 |
| **Vowels (%)** |  |  |  |  |  |  |  |  |  |
| Back rounded |  |  | 100 | 50 | 30 | 10 | 90 | 80 | 30 |
| Front unrounded |  |  |  | 50 | 70 | 90 | 10 | 20 | 70 |
|  |  | High dominance | Pointed | Light | Soft | Calming | Good | Bright | Big |
| **Consonants (%)** |  |  |  |  |  |  |  |  |  |
| Sonorants |  |  |  | 70 | 60 | 100 | 90 |  |  |
| Stops | Voiced |  |  |  |  |  | 10 |  | 20 |
|  | Unvoiced |  | 90 |  |  |  |  | 80 |  |
| Af/fricatives | Voiced |  |  |  |  |  |  | 10 | 60 |
|  | Unvoiced |  | 10 | 30 | 40 |  |  | 10 | 20 |
| **Vowels (%)** |  |  |  |  |  |  |  |  |  |
| Back rounded |  |  |  | 20 | 50 | 100 | 10 |  | 30 |
| Front unrounded |  |  | 100 | 80 | 50 |  | 90 | 100 | 70 |
